# Deep Learning-based Modeling Enhances Efficacy of Natural Ligand CAR Binders Targeting CD70

**DOI:** 10.64898/2026.09.06.749651

**Authors:** Amrik S Kang, Radhika Dalal, Mingcheng Li, Anupam Anand Ojha, Sujata Walunj, Caitlin E Cornell, Ruyan Rahnama, Abhilash Barpanda, Sanjana Prudhvi, Trishna S Patel, Mark Aspinall-O’Dea, Beth Walker, Clare Adams, Veronica Steri, Paul Phojanakong, Juan Antonio Camara Serrano, Hua Li, Yan Zeng, Jose Rivera, Daniel A Fletcher, Benjamin J Huang, Sonya M Hanson, Pilar Cossio, Tanja Kortemme, Justin Eyquem, Arun P Wiita

## Abstract

CD70 is well-recognized as a promising “pan-cancer” chimeric antigen receptor (CAR) T-cell target. Prior work has shown that a “natural ligand” (NL)-based CAR targeting CD70, employing its physiological interaction partner CD27, may have therapeutic advantages over antibody-based CARs. Yet while antibody-based CARs are routinely optimized by affinity maturation of their scFv, whether the binding sequence of an NL CAR can be engineered to improve its function remains unexplored. Here, we combined deep learning with physics-based modeling to redesign residues at the CD27:CD70 interface, identifying a CD27 variant, “N88A”, which enhances the efficacy of CD70-targeting CAR T-cells across models of acute myeloid leukemia, multiple myeloma, and renal cell carcinoma. Biophysical approaches, including molecular dynamics simulations, support a mechanism of increased binder conformational freedom underlying potency enhancement. Our work presents CD27_N88A_ CAR T-cells as a promising new therapeutic option and proposes that computational modeling could be applied to enhance efficacy of other NL-based immunotherapies.

## Introduction

Chimeric antigen receptor T-cells (CAR-Ts) are well-established as a breakthrough therapeutic technology for the treatment of multiple hematologic cancers^1–4^. The CAR design includes an extracellular domain for engaging a target antigen fused to intracellular T-cell signaling machinery^5,6^. The extracellular domain is often based on an antibody fragment such as a single-chain variable fragment (scFv); all currently commercially-available CAR-T therapies incorporate such antibody fragments^7^. However, CARs incorporating the endogenous protein binding partner to engage the target antigen have also been explored in the field, with several under clinical investigation^8–11^. These “natural ligand” (NL) CAR binders are noted to have potential advantages compared to antibody fragment counterparts, including a lower likelihood of receptor immunogenicity, greater surface receptor stability, and altered binding affinities, which in turn can reduce CAR-T exhaustion, improve CAR functionality, and reduce off-target binding^12–14^. Given the immense clinical need for new and improved CAR-T therapies for the numerous tumor indications that currently lack approved therapies, novel methods to identify and optimize NL CAR binders would be highly desirable.

Historically, most CAR binders have been based on scFvs originally derived from monoclonal antibodies raised against the antigen of interest^7^. Antibody discovery campaigns typically focus on the highest-affinity binders due to an assumption that higher affinity for target antigens would lead to improved CAR-T efficacy^15–17^. However, later studies have shown that this approach can lead to suboptimal CAR-T performance compared to moderate affinity binders, which often display lower exhaustion, improved expansion persistence, and even improved tumor discrimination from healthy tissues^18–20^. Improving scFv-based CARs by affinity tuning is well established^21–23^, in large part because antibodies provide an obvious target for mutagenesis: binding is concentrated in the complementarity-determining regions (CDRs), and CDR3 in particular offers a defined, tolerant locus for diversification. Natural ligands present no such roadmap. Their binding surfaces are distributed across a folded interface that also maintains the protein’s stability, and consequently NL CARs have rarely been engineered for improved function. In the sparse examples in the literature, changes in NL binder affinity have been made based on previously known receptor mutants^24^ or within the hinge and transmembrane region instead of the binding interface itself^25^. Notably, some NL binders have available high-resolution structures in complex with their targets^26,27^, providing an exceptional opportunity to fine-tune CAR affinities based on the alteration of known receptor-ligand contact points. Furthermore, NL CARs tend to have notably lower affinities to their target antigens compared to scFvs but often show comparable or even superior efficacy^14,28^, suggesting that the set of guidelines that have previously been compiled for CAR affinity tuning may not hold for NL CARs. This indicates a gap in the literature by describing a systematic approach to NL CAR optimization, particularly in the ability to identify potentially superior NL variants that are not already present in nature.

An excellent candidate system for a computational optimization campaign of an NL CAR is the CD27-CD70 receptor-ligand pair. CD70 is a member of the TNF ligand superfamily^27^ and has been identified as a cancer-specific target antigen in many cancers, including acute myeloid leukemia (AML)^29,30^, multiple myeloma^31^, B-cell cancers^32^, T-cell cancers^33,34^, and several solid tumors, such as renal cell carcinoma (RCC)^35^ and glioblastoma^36^. CD70 is absent from nearly all healthy tissues with the exception of transient expression on activated B-cells, T-cells, and dendritic cells, and has only one known endogenous receptor, CD27^37,38^. Prior work has shown significant promise in the use of CD27 to create a natural ligand CAR against CD70, with notable responses for some tumor models^25,39^, but relapse or incomplete response in others^40^, suggesting the potential for further optimization. This target and CAR system has been of special interest as a potential therapeutic in AML, which is the most common acute leukemia in adults and has a five-year survival rate under 30%^41^. Although several prior AML CAR-Ts have failed to progress through clinical trials, often due to high degrees of toxicity against hematopoietic stem and progenitor cell (HSPC) populations^42–44^, anti-CD70 therapies have shown much better safety profiles and promising preclinical and early clinical results^25,45^.

Here, we design and apply a computational approach, combining deep learning with physics-based modeling, to optimize an anti-CD70 CAR built on the extracellular domain of CD27. We identify mutational hotspots on the surface of CD27 and generate a library of CAR-T mutational variants to test their antitumor efficacy. In *in vitro* and *in vivo* assays, we identify a unique single point mutation in CD27, N88A, that displays improved tumor killing, *in vivo* expansion, and cytokine production. We utilize molecular dynamics combined with experimental studies to develop a biophysics-based mechanism of improved activity. Finally, we show that this CD27 mutant CAR maintains an excellent safety profile, with no off-target binding and no impacts on HSPC populations, supporting its further development as a potential best-in-class anti-CD70 CAR-T therapy. Furthermore, we propose that the approaches used here, leveraging advances in protein design and computational modeling, could be more broadly applied to enhance efficacy of other NL CARs.

## Results

### Deep learning and physics-based algorithms identify CD27 mutations with predicted altered energetics of CD70 binding

We developed a computational approach to generate new CD27-based binder variants for evaluation in CAR-T efficacy. We aimed to leverage recent advances in deep learning and neural networks for protein structural prediction and design^46–48^. This approach can be exceptionally valuable both in reducing the total number of receptor variants that need to be screened and in designing variants that contain multiple synergistic or codependent mutations, which would otherwise require testing an infeasibly large number of variants^49,50^. We therefore paired deep learning-based sequence design using ProteinMPNN^51^ with physics-based design and filtering using Rosetta FastDesign^52^. We first applied ProteinMPNN to the extracellular domain (ECD) of CD27, sampling the 24 residues that contact CD70 in the published crystal structure of the complex^53^ (**Fig. 1A**, *left*). ProteinMPNN designs contained between 2 and 8 different candidate residues at 17 of these positions (**Fig. 1A**, *middle*) which we used to constrain Rosetta interface design. We then used these proteinMPNN-generated residue choices to guide Rosetta interface design (**Fig. 1A**; see Methods). Filtering on predicted interface stability by Rosetta flex ddG^54^ and on shape complementarity relative to wild-type CD27-CD70 interface yielded 12 combinatorial designs, from which we selected four carrying between 2 and 5 mutations each (**Fig. 1B, S1A**).

**Figure 1.**
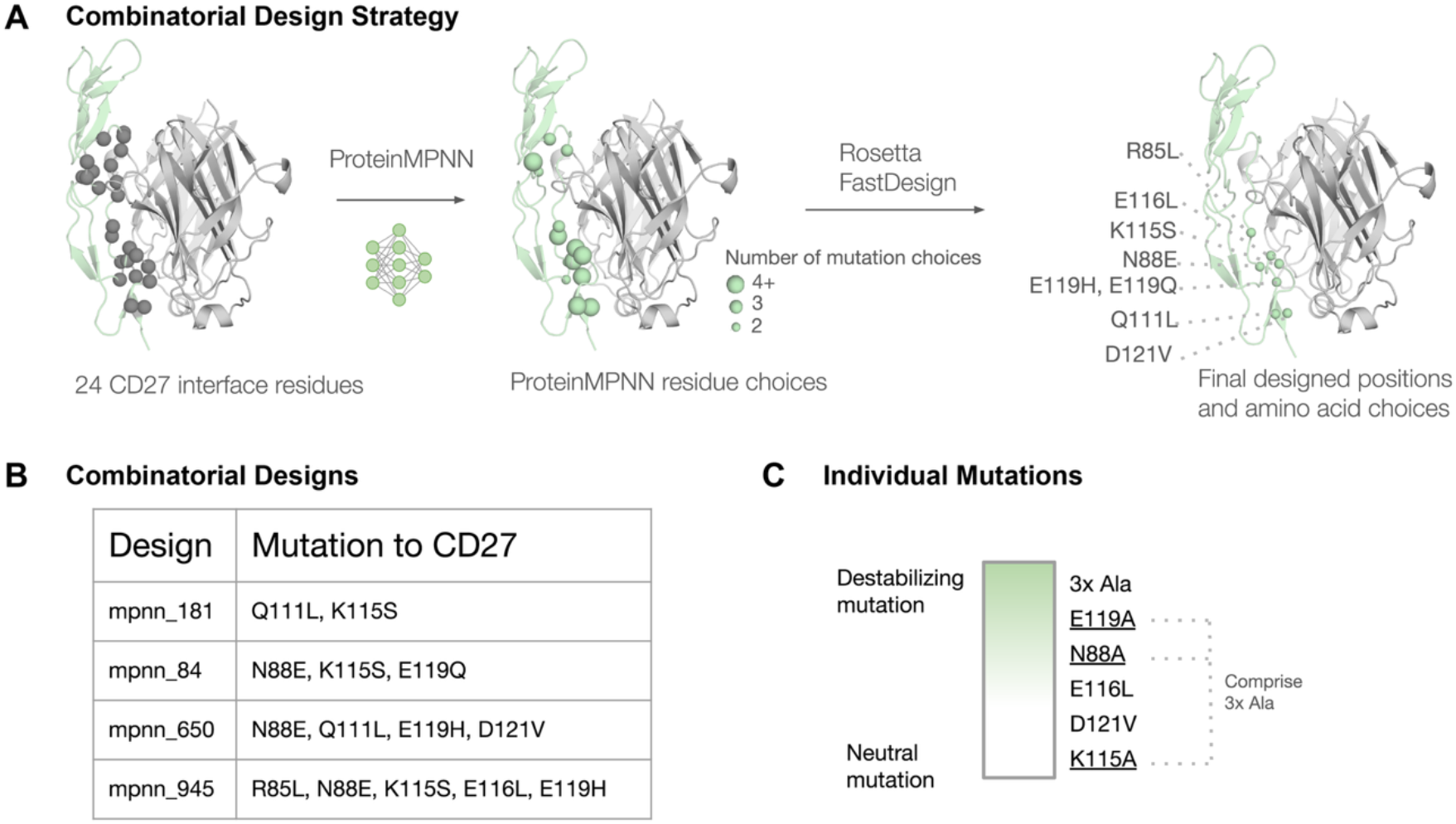
A design approach guided by deep learning and Rosetta sequence preferences. A. To initially discover combinatorial mutations at the CD27:CD70 interface, we first redesigned interface positions (gray spheres) using proteinMPNN. Mutation choices in MPNN designs at each position were then provided to Rosetta FastDesign. Designs were filtered that were either neutral or favorable in key Rosetta metrics (see Methods). Sample design positions shown as in green spheres. **B.** Sequence positions and mutations selected from the design strategy in A. were evaluated in specific design combinations. **C**. Additional individual hotspot interface interactions that were chosen for further evaluation. Heatmap indicates whether mutations were predicted to be stabilizing or neutral based on Rosetta energy score (see Methods).

We next decomposed these designs into their constituent single substitutions to determine which individual contacts drive the predicted changes in binding (**Fig. 1A**, *right*). For each mutation, we used Rosetta flex ddG to predict 1) the change in binding energy for the single mutation and 2) the change in binding energy for a mutation to alanine at the designated position (**Fig. 1C, S1B**). For initial *in silico* analysis, we selected a total of 8 single amino acid substitutions (R85L, N88E, Q111L, D121V, E119H, E116L, K115S, E119Q, **Fig. S1C**) in addition to the 4 combinatorial substitution designs described above (**Fig. 1C**). Optimization for tighter binding was not our only hypothesis; prior work on scFv-based CARs has shown that reduced-affinity binders can outperform their high-affinity counterparts^18,21,22^. We therefore included three variants predicted to destabilize the interface (N88A, E119A, 3-Ala) as well as one predicted neutral mutation (K115A) (**Fig. 1C, S1C**). Together, these variants provide a broad range of alterations for assessment by probing the most energetically significant amino acid contacts comprising the CD27:CD70 interface.

### In vitro assays identify promising CD27_var_ CAR-T designs

To assess the efficacy of our identified CD27_var_ designs as CAR binders, we cloned a subset of those identified from computational analysis into an AAV6 vector-compatible CAR cassette that can deliver a homology-directed repair template to insert the CAR into the *TRAC* locus^55^, allowing for uniform expression of each CD27_var_ CAR for direct comparison (**Fig. 2A**). We observed efficient *TRAC* knockout and CAR expression in 2 independent T-cell donors, with very similar knock-in efficiencies and homogeneous cassette expression levels across binder variants (**Fig. S2A-B**). To prevent cross-activation or fratricide, *CD70* was also knocked-out via CRISPR-Cas9 editing, as previously described for CAR T-cells against this target^31,56,57^. This set of CD27_var_ CAR-Ts was then stained with soluble recombinant CD70 conjugated to a fluorophore to allow for flow cytometry-based analysis of CD70 binding. We observed a distribution of relative CD70 binding, with most constructs staining a lower intensity compared to the CD27_WT_ binder but a small subset with comparable or slightly increased staining intensity (**Fig. 2B**). This result suggests that CD27_var_ binders can impact either the binding affinity, surface expression, or both, on CD70-targeting CARs.

**Figure 2.**
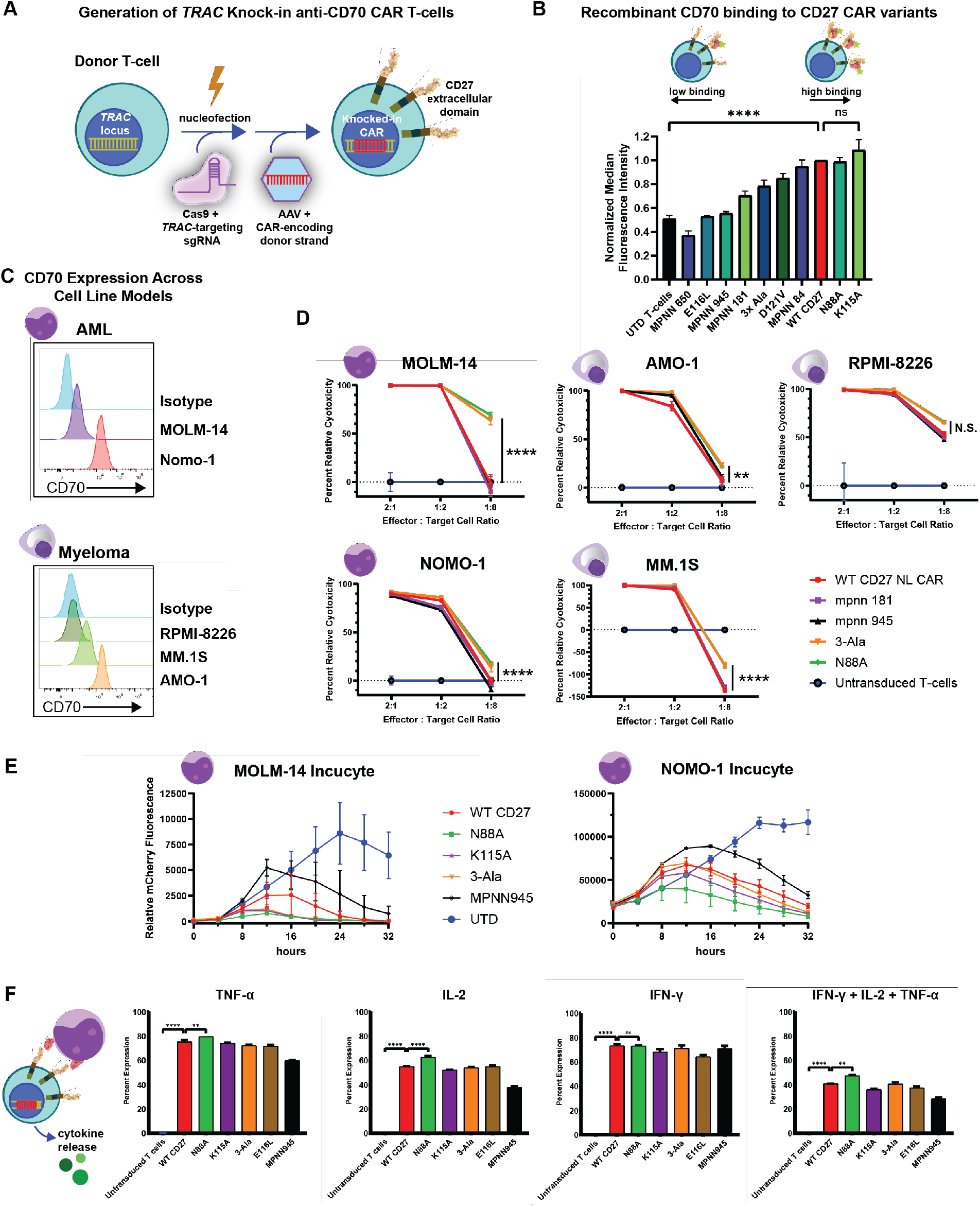
CD27_var_ CAR-Ts are expressed at the *TRAC* locus and exhibit differential tumor cytotoxicity. **A.** Cartoon depiction of AAV-based homology-directed repair method to express CD27-based, CD70-targeting CARs at *TRAC* locus. **B.** Binding to recombinant CD70 labeled with fluorescent marker by flow cytometry in *n*=2 T-cell donors normalized to CD27_WT_, error bars represent standard deviation. **C.** Expression of CD70 on multiple AML and multiple myeloma cell lines, as measured by flow cytometry. **D.** Relative percent cytotoxicity of CD27_var_ CAR-Ts in AML and multiple myeloma cell lines. Representative data from one donor shown for each CAR-T and AML target co-culture condition, *n*=2 T-cell donors tested. Statistical significance compares average cytotoxicity for CD27_WT_ vs CD27_N88A_ at the 1:8 E:T ratio using 2-way ANOVA and Tukey’s multiple comparisons test. **E.** Relative AML target cell killing in a 24- to 48-hour Incucyte live-imaging assay with CD27_var_ CAR-Ts in *n*=3 technical replicates, as measured by target cell mCherry fluorescence over time. Error bars represent S.E.M. **F.** Percent of CD27_var_ T-cells expressing TNF-α, IFN-γ, IL-2, and all 3 cytokines concurrently after a 6-hour co-culture with *n*=2 target AML cell conditions, as measured by intracellular flow cytometry. Statistical significance tested by one-way ANOVA and Dunnett’s multiple comparisons test, error bars represent S.E.M. *p<0.05; **p<0.01; ***p<0.005; ****p<0.001.

We chose 4 representative CD27_var_ designs, ranging across the distribution of binding affinities in **Fig. 2B**, to test CAR T-cell cytotoxic activity against different tumor types and antigen densities. We first evaluated CD70 expression on several representative acute myeloid leukemia (AML) and multiple myeloma (MM) cell lines, as both malignancies have previously been shown to be selectively targetable via CD70^14,30,31^. All cell lines tested were positive for CD70 expression, though with significant variation observed in relative expression levels (**Fig. 2C**). We then tested the ability for a subset of CD27_var_ CARs to kill each CD70-expressing tumor line in a 24-hour luciferase-based cytotoxicity assay, observing that most variants performed quite similarly to the CD27_WT_ CAR, but that the CD27_N88A_ and CD27_3-Ala_ CAR-Ts displayed subtle yet consistent improvements in CAR-T killing ability (**Fig. 2D**). To further validate these results and better characterize the cytotoxicity kinetics of these CAR-T variants, we performed a live cell imaging-based co-culture experiment with the AML cell lines MOLM-14 and NOMO-1, observing that the CD27_N88A_ variant consistently cleared the tumor cells most rapidly, with significant improvement compared to the CD27_WT_ CAR-T (**Fig. 2E**).

Finally, we evaluated if these differences in *in vitro* tumor cytotoxicity were correlated with T-cell cytokine secretion in a short-term stimulation assay. In a 6-hour tumor co-culture experiment, CD27_N88A_ CAR was observed to have a larger percent of T-cells that secreted TNF-α, IFN-γ, and IL-2 compared to CD27_WT_ CAR (**Fig. 2F**). Conversely, CD27_mpnn945_, which had previously been seen to underperform CD27_WT_ in cytotoxicity assays, also had a smaller percent of T-cells that were triple-positive for these cytokines upon tumor stimulation, suggesting that the differences in cytotoxicity were correlated with short-term cytokine production.

### CD27_N88A_ CAR outperforms the CD27_WT_ CAR in vivo in AML and MM models

Encouraged by the improved efficacy we observed in several of our CD27_var_ CARs *in vitro*, we tested a subset of variants in an aggressive *in vivo* cell line model of AML. We first infused 1e6 MOLM-14 cells into NSG mice, followed by 5e5 CD27_WT_, CD27_var_, or untransduced CAR-T cells 4 days later, monitoring tumor growth by bioluminescence imaging (**Fig. 3A**). Notably, mice treated with CD27_WT_ CAR could not fully control this aggressive tumor model, with a median survival in this group of 38 days, compared to 21 days in the negative control untransduced T-cell group. In comparison, the CD27_N88A_, CD27_K115A_, and CD27_MPNN945_ groups all showed reduced tumor burden and improved survival compared to the CD27_WT_ group, with all mice in the CD27_N88A_ group maintaining no detectable tumor until the conclusion of the study 85 days after CAR-T treatment (**Fig. 3B-C**). This dramatic improvement was correlated with improved CAR-T expansion and persistence in the treated mice, as measured by peripheral blood expansion, with the CD27_N88A_ group showing the best sustained persistence of CAR-Ts (**Fig. 3D**). Interestingly, CD27_MPNN945_ showed improved expansion and persistence as well, likely contributing to the improved survival compared to CD27_WT_, despite it notably worse performance in our *in vitro* assays.

**Figure 3.**
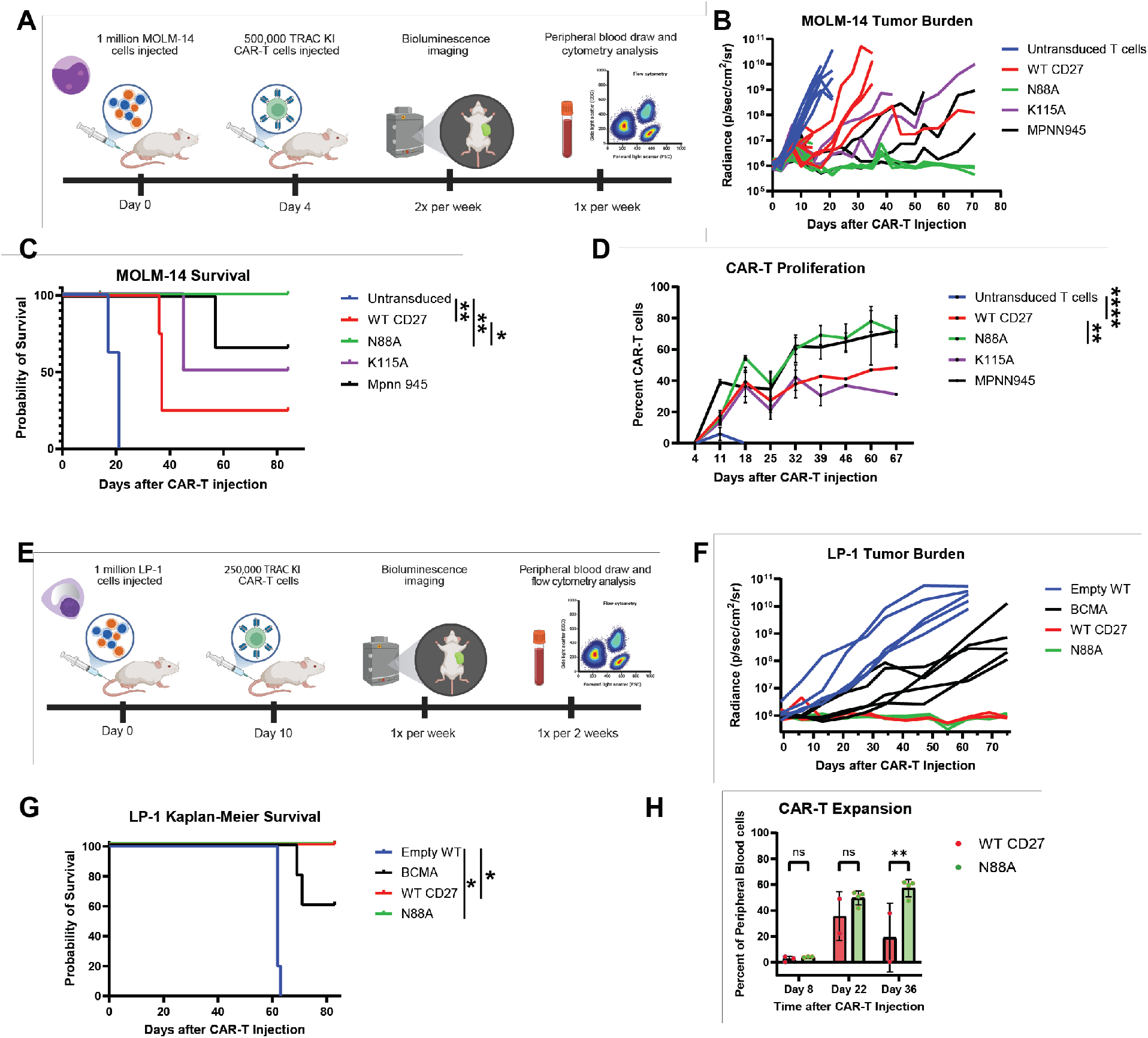
CD27_N88A_ CAR T-Cells achieve superior tumor control in *in vivo* models of AML and multiple myeloma. **A.** Cartoon overview timeline of *in vivo* CAR-T mouse experiment with MOLM-14 cell line. **B**. Quantification of tumor burden following bioluminescence imaging of mice after CAR-T injection. **C.** Kaplan-Meier survival curve by treatment group, statistical significance determined by Mantel-Cox test. **D.** Flow cytometry-based measurement of CAR-T proliferation in each treatment group as a percent of all live non-RBC events from peripheral blood draws. Statistical significance determined by 2-way ANOVA and Tukey’s multiple comparisons test. **E.** Cartoon overview timeline of *in vivo* CAR-T mouse experiment with LP-1 cell line. **F.** Quantification of tumor burden following bioluminescence imaging of mice after CAR-T injection. **G.** Kaplan-Meier survival curve by treatment group, statistical significance determined by Mantel-Cox test. **H.** Flow cytometry-based measurement of CAR-T proliferation in each treatment group as a percent of all live non-RBC events from peripheral blood draws, statistical significance determined by 2-way ANOVA and Šídák’s multiple comparisons test. *p<0.05; **p<0.01; ***p<0.005; ****p<0.001.

We next tested the efficacy of our lead candidate CD27_N88A_ CAR against a tumor model for high-risk MM, which we recently identified as a promising cancer indication for anti-CD70 CAR-T therapy^31^. We utilized the LP-1 cell line, which harbors a t(4;14) translocation representative of a genomically-defined subtype of high-risk MM. We infused 1e6 LP-1 cells into NSG mice, followed by 2.5e5 CD27_WT_ or CD27_N88A_ CARs generated by *TRAC* knock-in, compared to 5e6 anti-BCMA (mimic of FDA-approved ciltacabtagene autoleucel; generated by lentivirus reflective of the clinical manufacturing process) (**Fig. 3E**). Remarkably, despite the 20-fold lower CAR-T dose for the anti-CD70 CAR-Ts, we saw that both CD27-based CAR conditions outperformed the anti-BCMA CAR – the current gold-standard CAR-T therapy for MM – with no detectable tumor in the CD27 CAR-treated mice at the conclusion of the study (**Fig. 3F-G**). In concordance with the MOLM-14 *in vivo* study, CD27_N88A_-treated mice saw greater CAR-T proliferation in the peripheral blood compared to the CD27_WT_ group (**Fig. 3H**). These results highlight the improved *in vivo* performance of the CD27_N88A_ construct in multiple hematological tumor models and its superiority against a clinically-approved CAR construct.

### CD27_N88A_ CAR displays an altered transcriptional state against CD70-expressing tumors in vivo

As there are a number of key signaling pathways for CAR-T activation^58,59^, we next explored how transcriptional changes between CD27 variants could help drive the observed differences in CAR-T function. We performed an *in vivo* tumor challenge experiment using the MOLM-14 cell line, in a design similar to that in **Fig. 3A**, treating the mice with either CD27_WT_ or CD27_N88A_ CARs. Ten days after CAR-T infusion, mice were sacrificed and their spleens and bone marrows were harvested to capture CAR-Ts, which were then isolated via FACS sorting (**Fig. S3A**). As before, we observed a significantly higher level of CAR-T proliferation in mice treated with CD27_N88A_ CARs compared to CD27_WT_ CARs (**Fig. S3B**). We then performed single-cell RNA-sequencing to compare the cell states and transcriptional profiles of CAR-T cells in these two treatment groups. Notably, the CD27_N88A_ group expressed a statistically significantly higher level of *FosB* (**Fig. S3C**), a subunit of the AP-1 transcription factor that has been well-characterized to drive T-cell differentiation and effector function following activation^60,61^.

Furthermore, within the CD4 subset of CAR-Ts, which had previously been observed to be the primary contributor to CD27 CAR-T expansion, *MKI67* levels were observed to be moderately higher in CD27_N88A_ CARs, consistent with the improved expansion phenotype (**Fig. S3D**). Taken together, these results support the therapeutic potential of the CD27_N88A_ variant.

### Biophysical analyses show that CD27_N88A_ binds CD70 with greater affinity and altered kinetics compared to CD27_WT_

Given the phenotypic results above, we explored whether biophysical characteristics of the N88A mutant may underlie improved CAR efficacy. We first recombinantly expressed extracellular domains from the CD27_N88A_ and CD27_WT_ receptors and performed biolayer interferometry against immobilized trimeric CD70 ligand (**Fig. 4A**). Unexpectedly based on our initial predictions (**Fig. 1C**), CD27_N88A_ was observed to have an approximately two-fold higher affinity for CD70 compared to CD27_WT_, at 19.36 ± 0.10 nM vs 36.94 ± 0.12 nM, respectively.

**Figure 4.**
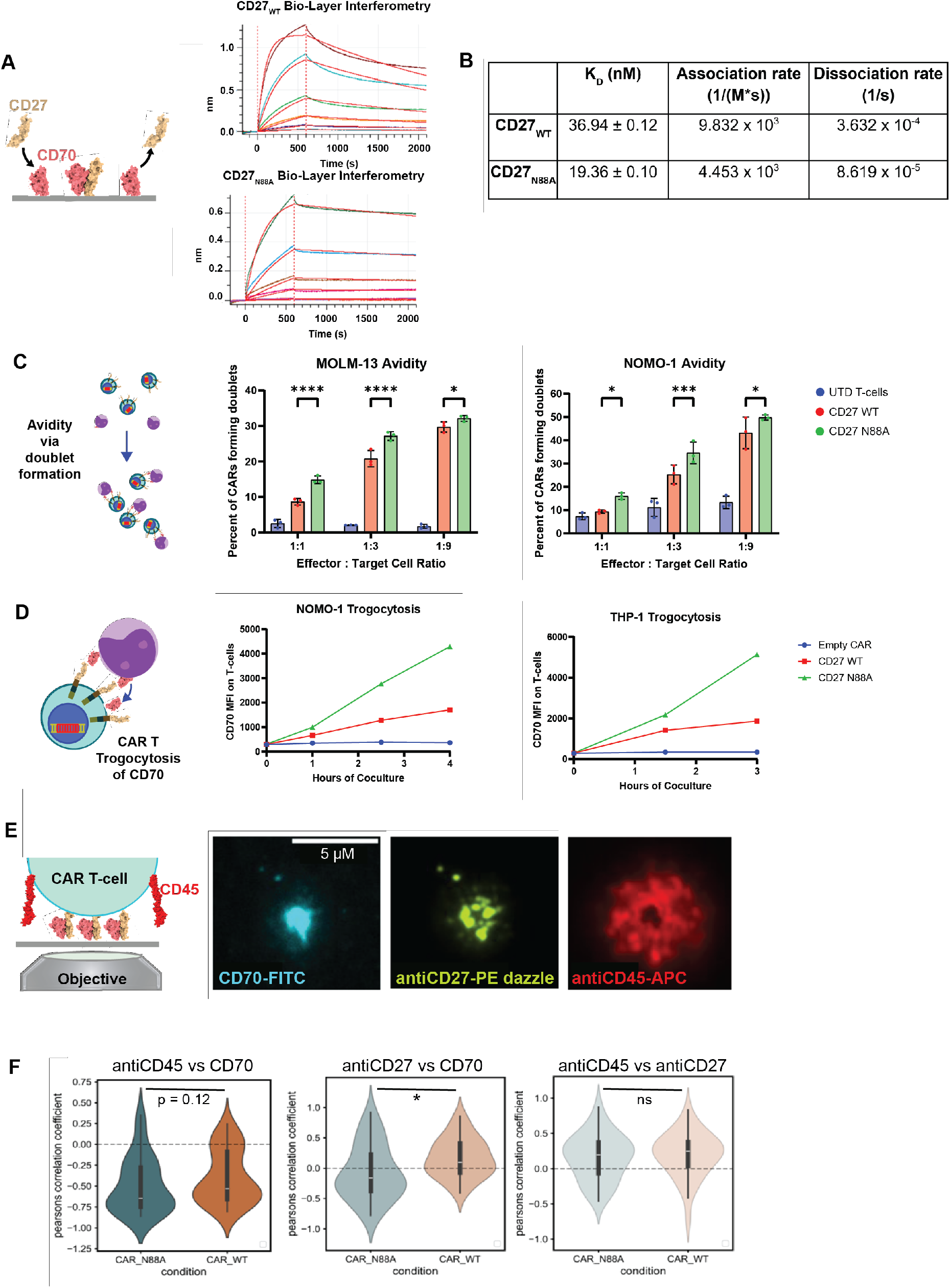
Biophysical characterization and fluorescent microscopy reveal that CD27_N88A_ has improved binding to CD70 and leads to altered CAR-T synapse morphology. **A.** Association and dissociation best-fit curves of CD27-based recombinant protein to sensor-immobilized CD70 via biolayer interferometry, measured at 1080, 360, 120, 40, 13.3, and 4.4nM of CD27_var_ protein concentration. **B.** Biolayer interferometry best fit-derived dissociation constant, association rate, and dissociation rate for recombinant CD27_WT_ or CD27_N88A_ binding to recombinant CD70. **C.** Relative avidity of CD27-based CARs, measured as a percentage of CAR-Ts forming doublets with AML target cells by flow cytometry. n=3 technical replicates, *p*-value by 2-way ANOVA with two-stage step-up FDR. **D.** Quantification of trogocytosis causing expression of CD70 on the surface of CAR-T cells following co-incubation with CD70-expressing AML cell lines NOMO-1 and THP-1. **E.** Representative images of total internal reflection fluorescence microscopy of CD27_N88A_ CAR co-incubated with CD70 displayed on lipid bilayer. **F.** Violin plots of the Pearson correlation between the relative intensity of fluorescent markers in n=34 images of each CAR variant synapse; statistical significance determined by unpaired t-test. *p<0.05; **p<0.01; ***p<0.005; ****p<0.001

This improvement was observed to be driven primarily by a more than four-fold slower dissociation rate for CD27_N88A_, whereas the association rate is approximately two-fold slower compared to CD27_WT_ (**Fig. 4B**). These results suggest that the CD27_N88A_ CAR may take longer to bind the CD70 complex, but that the resulting interaction is more stable than with the CD27_WT_ binder. These changes in interaction energetics and kinetics may potentially relate to the altered signaling and transcriptional programs observed in T cells expressing this CAR variant.

We next tested if these differences in individual CAR affinity also extended to CAR avidity. We used an established flow cytometry-based CAR-target cell doublet assay to approximate the relative avidity of our CD27-based CARs^62,63^. When co-incubated with the MOLM-13 and NOMO-1 cell lines, CD27_N88A_ CAR-Ts formed significantly more doublets with target cells than CD27_WT_ CAR-Ts, indicating that the cumulative avidity of CAR interactions was greater in our lead candidate CAR construct (**Fig. 4C**).

Receptor-ligand affinity has previously been shown to be correlated with trogocytosis, a mechanism by which cells can acquire molecules expressed on other cells with which they come in direct contact^64^. In a short-term assay, we observed marked time-dependent trogocytosis of CD70 antigen onto CAR T-cells when co-cultured with the AML-cell lines NOMO-1 and THP-1, with notably higher levels of CD70 trogocytosis observed in the CD70_N88A_ CAR compared to the CD70_WT_ CAR (**Fig. 4D**).

Intrigued by these observed improvements in CD70 engagement for the CD27_N88A_ construct, we sought to visualize these differences at the scale of the CAR synapse with target antigen. We used total internal reflection fluorescence (TIRF) microscopy to observe the clustering of CAR molecules and CD45 on the surface of CAR-T cells as they came in contact with fluorescently labeled target CD70 molecules displayed on a phospholipid bilayer (**Fig. 4E**). We observed that CD27_N88A_ CAR T-cells formed synapses that frequently differed from CD27_WT_ CARs. Most notably, the CD70 fluorescence showed significantly greater segregation from stained CD27 CAR for CD27_N88A_ compared to that of CD27_WT_ as measured by their Pearson correlation coefficient (**Fig. 4F**). These data are consistent with the tighter binding of CD27_N88A_ to CD70, causing the anti-CD27 antibodies to be excluded from the center of the CAR T-target interface and preferentially label peripheral or unbound CAR molecules. Together, these results highlight the major impact of a single point mutation on the protein-protein interaction at the CD27:CD70 interface.

### Molecular dynamics simulations predict that CD27_N88A_ forms a more dynamic and stable complex with CD70

Given the unexpected observation that the N88A mutation led to increased binding affinity for CD70, we turned to all-atom molecular dynamics simulations to model the structural and dynamic consequences of this mutation. Three independent replicates of 2.25 μs were run for each complex, with a cumulative sampling of 13.5 μs, starting from the CD27-CD70 crystal structure (PDB 7KX0), with all crystallographically resolved N-linked glycans retained. We first generated free energy landscapes of the CD27_N88A_ and CD27_WT_ binders in complex with CD70, projected onto two collective variables: the root mean square deviation (RMSD) with respect to the crystal structure and the radius of gyration (R_g_) of the C_α_ atoms of the trimeric CD27-CD70 functional units. A well-defined free energy minimum centered at RMSD ≈ 2.0 - 3.0 Å and R_g_ ≈ 26.5 Å was observed for CD27_WT_, with the population confined within this single basin across all replicates. In contrast, CD27_N88A_ explores a broader conformational space, with multiple metastable states sampled across a range of RMSD (2.5 - 5.0 Å) and R_g_ (26.0 - 27.5 Å) values (**Fig. 5A, S4A**). Multiple low energy basins indicate the ability of the mutant to adopt multiple stable conformations, while allowing for structural fluctuations without dissociation.

**Figure 5.**
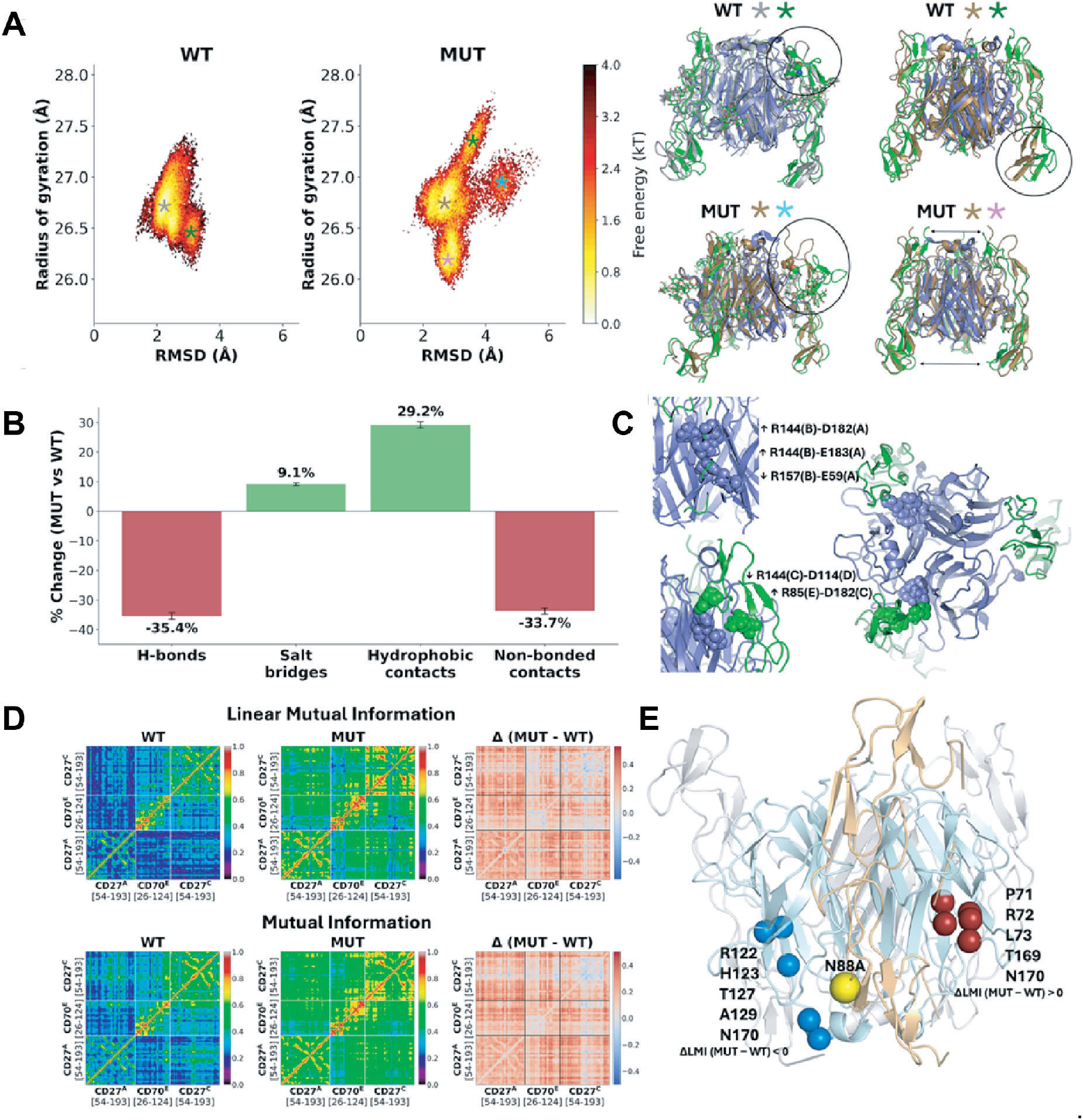
Molecular dynamics analysis of CD27-CD70 interface dynamics. **A.** Two-dimensional free energy landscapes for the WT and N88A complex are shown with representative conformational states marked by colored asterisks and corresponding structural overlays illustrating interface rearrangements. **B.** Percent change in interfacial interactions upon N88A mutation is shown, with error bars representing S.E.M. across three replicates. **C.** Structural mapping of salt bridge changes is displayed, where arrows denote increased (↑) or decreased (↓) occupancy in the N88A complex. **D.** Residue-residue correlation matrices were computed using linear mutual information (top) and mutual information (bottom). **E.** Eigenvector centrality changes are mapped onto the structure, where blue spheres indicate decreased centrality, red spheres indicate increased centrality, and the yellow sphere marks the N88A mutation site.

To understand the molecular basis for the enhanced conformational flexibility observed in the mutant dynamics within the free energy landscape, we next performed trajectory analyses around the N88A mutation sites in each CD27 monomer of the trimeric complex, averaged across three independent replicates. The most pronounced change was observed in the hydrogen bonding network (at a 3.5 Å cutoff), showing a 35.4% ± 1.1% reduction in hydrogen bonds (H-bonds) in the mutant (**Fig. 5B, S4B**), indicating the loss of intermolecular hydrogen bonds by asparagine amide groups. We observed the additional loss of several H-bonds with this mutation, but with a compensatory increase in several salt bridges (increased by 9.1% ± 0.5%) and the emergence of new electrostatic contacts (**Fig 5B-C, S4C-D, Table S1-2**).

Hydrophobic contacts (with 5.0 Å cutoff) also increased by 29.2% ± 1.1% (**Fig. 5B, Table S3**), likely due to the substitution of a polar asparagine with alanine, which introduces new hydrophobic surface area for interactions with neighboring residues. Total non-bonded contacts (with 4.5 Å cutoff) decreased by 33.7% ± 1.1% (**Fig. 5B**, **S4E, Table S4**), suggesting that the N88A interface is less densely packed than the WT interface. Fewer H-bonds and non-bonded contacts in the mutant are expected to weaken binding, and the measured slower dissociation rate suggests that the stability gained by the mutant is kinetic rather than a consequence of tighter packing, arising from the exchange of a small number of directional hydrogen bonds for a larger set of non-directional hydrophobic contacts that are maintained across multiple interface geometries.

Next, residue-residue correlation networks were computed using linear mutual information (LMI) and mutual information (MI) metrics to analyze both direct mechanical coupling and allosteric effects caused by the N88A mutation. CD27_N88A_ exhibited increased inter-domain correlations compared to the CD27_WT_. Mean LMI values were increased by 58% for CD27_N88A_ vs CD27_WT_, with a similar 47% increase for MI (**Fig. 5D**). The difference correlation maps (Δ = CD27_N88A_ - CD27_WT_) showed predominantly positive values across both metrics, with LMI differences exceeding +0.15 in multiple regions spanning the CD27-CD70 interface (**Fig. 5D**). This increase in correlated motion suggests that the CD27 and CD70 subunits move more coherently in the N88A complex, leading to improved interface reorganization. Eigenvector centrality analysis, which identifies communication hubs within the correlation network, revealed a significant redistribution of network topology upon mutation (**Fig. 5E, S4F-G**). In the WT complex, the highest-centrality residues clustered in the CD70 chain C (T169, I132, N170, D165, and R163, spanning residues 119 - 170), serving as the primary communication hub coordinating intermolecular dynamics. The N88A mutation induced a dramatic shift in this network architecture, where the centrality decreased substantially in this WT hub region. This rewiring of allosteric communication pathways may have significant functional consequences.

Taken together, these findings from molecular dynamics simulations provide a unifying biophysical explanation for our observations, whereby an expanded conformational ensemble suggests that the N88A mutation increases the flexibility of the trimeric complex while maintaining overall structural integrity. We interpret these results as evidence that the energetics of the N88A mutation, while slowing initial complex association, allow the binder to “breathe” with the target rather than dissociate, such that the escape requires exit from an ensemble of bound states rather than a single bound state. This is consistent with the longer dwell time, a slower dissociation rate, and, ultimately, higher affinity.

### CD27_N88A_ TRAC CAR T-cells show strong potency in a solid tumor model in vivo

In addition to its expression on AML, MM, and other hematologic malignancies, CD70 is expressed on numerous solid cancers, most notably including renal cell carcinoma (RCC). Immunotherapies developed against CD70 for these indications have shown promising preliminary results, although they have generally required high doses to be initially delivered to be effective^56,57,65^. We wanted to test if our CD27_N88A_ construct would also show improved performance in an RCC solid tumor model, thereby suggesting generalizable improved performance to all CD70-expressing malignancies.

We first confirmed the expression of CD70 on two RCC cell lines, 769-P and ACHN (**Fig. S5A**). We implanted 5e6 ACHN tumor cells subcutaneously into the flank of NSG mice, followed by a a low dose of 5e5 CD27_WT_, CD27_N88A_, or untransduced CAR T-cells 11 days later. Additionally, we performed a rechallenge of 5e6 ACHN tumor cells after the initial tumors were controlled into the opposite flank of the mice on day 29 (**Fig. S5B**). We observed significant tumor control by both CAR constructs, which was maintained following the tumor re-challenge on day 37, and a notable trend toward improved efficacy of CD27_N88A_ compared to CD27_WT_ (**Fig. S5C-E**).

### CD27_N88A_ shows no off-target toxicity and has no impact on hematopoietic stem and progenitor cell populations

One potential concern with generating mutations to alter the binding characteristics of a natural ligand-based binder is the potential for the new receptor to develop off-target binding and cause unintentional toxicity. To explore this possibility, we utilized the Retrogenix^®^

Cell Microarray screening platform to test for off-target binding. In this assay, recombinant CD27_N88A_ protein was tested for binding to 6108 unique human cell membrane and secreted proteins, including the positive control CD70 target antigen. We first assessed the on-target and background binding signals of CD27N88A against CD70, CD86, or EGFR. Adequate CD70-specific signals were observed at binder concentrations of 0.4, 1.0, and 4.0 μg/mL (**Fig. S6A**). Next, we tested CD27_N88A_ against the surface protein library in quadruplicate, observing the expected interaction with CD70. Interactions identified in the library screen, on fixed cells, were then replicated in a confirmation screen performed on live cells. Other than the intended interaction with CD70, the only other interactions were mediated by binding of FCGR3A to the human Fc (hFc) domain fused to CD27, and were also observed for the positive control CTLA4-hFc (**Fig. 6A, S6B**). Taken together, this broad-scale screening platform highlights that the N88A mutation in CD27 does not result in any off-target activity compared to the physiological WT sequence of CD27.

**Figure 6.**
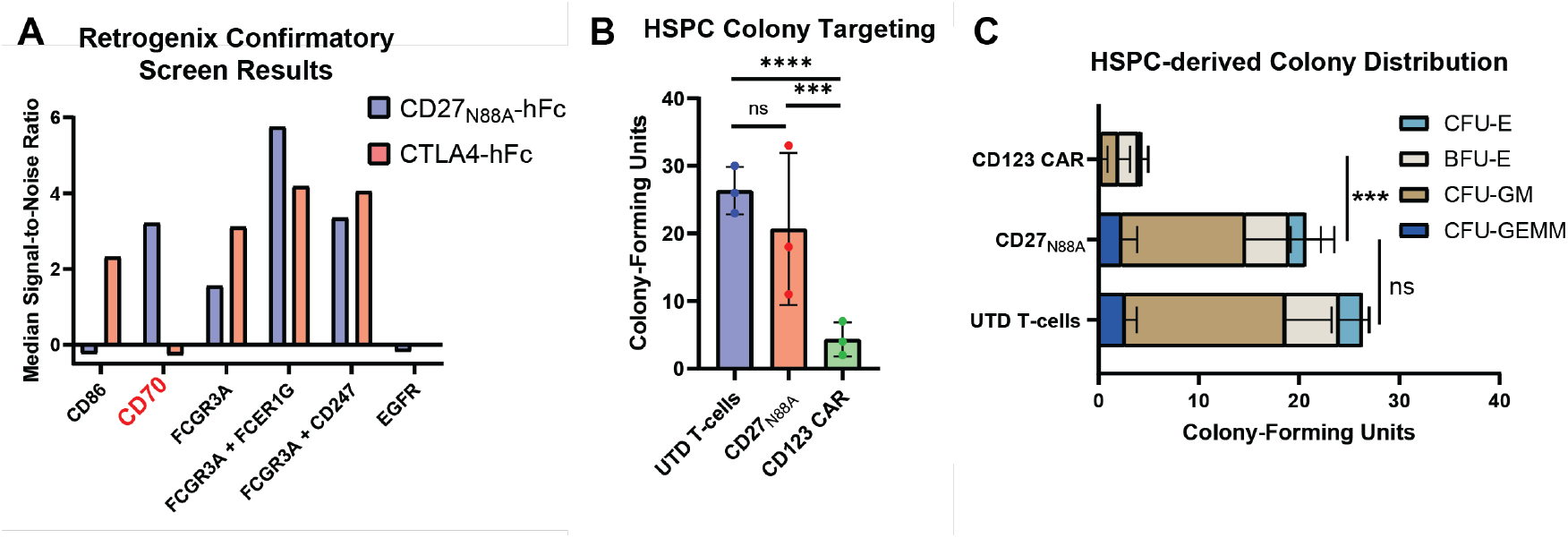
CD27_N88A_ variant does not lead to off-target toxicity. **A.** Representative quantitative results from Retrogenix screen confirmation that CD70_N88A_ variant solely interacts with target CD70 and CTLA4 positive control solely interacts with target CD86. As expected, fused Fc domain on both recombinant proteins interacts with Fc receptor domain FCGR3A. Neither protein interacts with EGFR nor any other protein on the screening panel. **B.** Total number of CD34+ HSPC-derived colonies counted in methylcellulose media following a co-incubation with N88A CAR or negative and positive control CAR-Ts. *n* = 3 technical replicates, *p*-value by one-way ANOVA with Tukey’s multiple comparisons test. **C.** Breakdown of types of HSPC-derived colonies based on morphology analysis as identified in panel B. UTD = untransduced. CFU = colony-forming unit. BFU = blast-forming unit. E = erythroid. GM = granulocyte/macrophage. GEMM = granulocyte, erythroid, macrophage, megakaryocyte. *p*-values indicative of total colonies, as in B. *p<0.05; **p<0.01; ***p<0.005; ****p<0.001.

As previously noted, one of the proposed benefits of targeting CD70 is its relative lack of expression on healthy tissues, particularly on HSPCs, which frequently express other AML antigen targets^30,66–68^. To confirm that the CD27_N88A_ mutation does not impact HSPCs, we performed a colony-forming assay, co-incubating CAR-Ts with CD34+ HSPCs from healthy donors (**Fig. 6B**). CD27_N88A_ CAR had no impact on HSPC colony growth compared to untransduced T-cells, whereas an anti-CD123 CAR significantly reduced the total number of viable HSPC colonies (**Fig. 6B**). Specifically, the greatest reduction in colonies was observed in the CFU-GM subset (**Fig. 6C**), as expected based on known CD123 expression on granulocyte and monocyte precursors^66,69^. These assays demonstrate the lack of off-target interactions and minimal on-target, off-tumor toxicity for CD27_N88A_ CARs, underscoring their strong potential for further clinical translation.

## Discussion

Our work demonstrates a novel approach to optimizing natural ligand-based (NL) CAR binders, applied here to the well-characterized tumor antigen CD70^30,56,70–72^. Utilizing protein design algorithms for affinity tuning has been previously reported for antibody fragment-based binders^73^, but to our knowledge has never been used for NL CARs. A limited set of literature does exist on improving NL-based CARs, but these have lacked a systematic and generalizable approach to binder optimization^24,25,62,74^. Indeed, given that NL CARs frequently have a greater number of individual interaction residues and generally weaker affinities compared to antibody-based binders^13,27^, they appear to be excellent targets for improvement by *in silico* prediction algorithms. Here, we show that although *in silico* predictions cannot alone identify the most superior binders, when paired with comprehensive assays to evaluate CAR-T function, they can allow for the rapid screening and identification of lead candidate NL CAR binders.

A significant body of literature has probed where the optimal binding affinity lies for CAR-T binders, although this has almost entirely been evaluated in antibody fragment-based binders, which generally have stronger affinities to their target antigens than natural ligand-based binders^13^. Although early attempts at optimizing binders generally worked to improve binder affinity^22,75^, more recent work has suggested the existence of an affinity “sweet spot”, as an overly strong receptor-ligand interaction may lead to faster T-cell exhaustion and lower processivity^20,69^. In a notable example, a CD19-targeting CAR developed using a significantly weaker affinity than the FDA-approved FMC63 clone-based CAR showed lower exhaustion markers and superior persistence in clinical trials^76^. Biochemical analysis revealed that this binder had a similar rate of association but a faster rate of dissociation, which in turn resulted in a higher K_D_ value^76^. In contrast, we show that the CD27_N88A_ binder, which outperforms the CD27_WT_ binder in a CAR format, has tighter binding to CD70, as measured by a lower K_D_ value, and that this is driven primarily by a significantly slower dissociation rate. We hypothesize that this may allow for prolonged T-cell synapse formation and downstream signaling, which in turn may lead to more effective effector function for the CAR T-cells. This is supported by our findings that CD27_N88A_ CAR-Ts produce more effector cytokines when stimulated by tumor cells and have altered CAR signaling and transcriptional states after incubation with CD70-expressing cell lines. Given the weaker interaction of CD27_WT_ with CD70 compared to most antibodies to their ligands, it may not be surprising that the “sweet spot” for this CAR interaction requires a stronger interaction than the wild-type receptor. Indeed, it may be true that similar optimizations of other NL CARs may also result in stronger interactions with their ligands, although more data will be needed to support this hypothesis.

Our analysis also includes a unique application of molecular dynamics simulations to propose a mechanistic model for the enhanced binding properties of the CD27_N88A_ complex. The asparagine-to-alanine substitution removes hydrogen bonding at the interface, replacing them with hydrophobic contacts. These new interactions initiate a cascade of predicted consequences, namely interface reorganization, enhanced conformational sampling, rewired allosteric communication, and slowed conformational kinetics. The net effect is a predicted transformation from a lock-and-key binding (WT) to a dynamic fit binding (N88A). In the WT complex, hydrogen bonds are predicted to adopt a single stable conformation, whereas in the mutant, hydrophobic contacts are predicted to maintain stability across multiple conformations as dissociation requires escape from an entire ensemble of bound states. This model of dynamic stability with conformational flexibility enables multiple conformational states and provides a molecular framework for understanding how specific mutations can improve natural ligand-based CAR binders and may inform future optimization strategies for this class of therapeutics.

We also note that our lead candidate CD27_N88A_ CAR, expressed as a gene knock-in under the control of the *TRAC* locus, likely represents a best-in-class CD70-targeting CAR-T therapeutic. Prior reported CD70-targeting CARs have required dosages between 1 and 5 million CAR T-cells to treat cell line mouse xenograft models of hematological cancers^14,25,77^, and between 1 and 10 million CAR T-cells to treat solid cancer mouse xenograft models^56,57^. In comparison, we show here persistent, curative responses with 500,000 CAR T-cells for aggressive models of acute myeloid leukemia and renal cell carcinoma, and with only 250,000 CAR T-cells for a high-risk molecular subtype of multiple myeloma. Notably, this latter experiment showed significantly better control than 5 million CAR T-cells expressing a cilta-cel^78^ mimic targeting BCMA, which is considered to be the gold standard cell therapy for multiple myeloma^79^. The extraordinarily robust antitumor efficacy and notable expansion and persistence of the CD27_N88A_ CAR in multiple tumor indications, in combination with its lack of observable off-target effect or toxicity against HSPCs, provides a strong rationale for its continued development as a potential therapeutic for future translation to the clinic. Future work will evaluate whether the CD27_N88A_ binder may further improve the more complex HLA-independent T-cell (HIT) engineered cellular therapy strategy; this approach, incorporating an anti-CD70 antibody fragment, recently showed potential to enhance T-cell sensitivity to low CD70 antigen density on tumors^80^.

We also acknowledge several limitations in our study. Notably, there was not a strong correlation between the stability of the CD27_var_-CD70 complex as predicted by Rosetta and the variants that were found to be the most efficacious in functional studies. This finding suggests that while deep learning approaches were able to identify critical residues that would impact CAR efficacy when mutated, *in silico*-predicted receptor-ligand complex stability may be an incomplete metric for predicting the efficacy of a CAR binder. Future work will include screening approaches to evaluate a much larger set of computationally predicted CD27 mutants, generating a dataset that could be used to train more accurate models to predict CAR functionality. Furthermore, a large body of literature has shown that hinge, transmembrane, costimulatory, and intracellular signaling regions of CARs play significant roles in the cytotoxic ability, *in vivo* efficacy, and overall behavior of CAR T-cells^25,81–83^. In this study, all CD27_var_ CARs were tested using a single transmembrane and intracellular signaling framework. Future work will systematically evaluate the optimal non-binder CAR design for the CD27_N88A_ CAR, which will also serve as an important step toward clinical translation.

Overall, this work illustrates a deep learning-based approach to an optimized CD27-based CAR. We further established biophysics-based mechanistic insight into the reasons for its improved functionality of the CD27_N88A_ mutant, including improved CAR signaling, tighter binding affinity, and altered dynamics that allow for greater stability at the CAR synapse. Our work broadly illustrates that for development of natural ligand CARs, it is indeed possible to improve on Nature’s wild-type ligand sequences through computationally driven protein engineering strategies. Furthermore, these results strongly support additional development of this specific cell therapy design towards clinical translation in CD70-positive malignancies.

## Author Contributions

A.S.K., M.L., S.W., C.C., R.R., A.B., S.P., T.P., M. A.-O., B.W., C.A. performed in vitro experiments and analyzed data. R.D. performed protein design simulations and Rosetta analysis. A.A.O., S.H., and P.C. performed molecular dynamics simulations. B.J.H. analyzed RNA sequencing data. V.S., P.P., J.A.C.S., H.L., and J.S. performed murine studies. A.P.W., J.E., and T.K. acquired funding and supervised the work. A.S.K. and A.P.W. wrote the manuscript with additional contributions and approval by all authors.

## Acknowledgements

This work was supported by the Myeloma Solutions Fund, a Translational Research Program award from Blood Cancer United, CRISPR Cures for Cancer, the Silicon Valley Community Foundation, the Cookies for Kids Cancer Foundation and the UCSF Multiple Myeloma Translational Initiative (all to A.P.W.). The Eyquem laboratory is grateful for the support from the Parker Institute for Cancer Immunotherapy, the Pew Charitable Trust and The Alexander and Margaret Stewart Trust, the Grand Multiple Myeloma Translational Initiative, the Multiple Myeloma Research Foundation, the Baszucki Lymphoma Fund, CRISPR Cures for Cancer and the Weill Cancer Hub West. During completion of this work A.S.K. was supported by NIH F30CA298678 and T32GM141323. R.D. was supported by the National Science Foundation Graduate Research Fellowship Program. R.R. is supported by NIH K12HD105250-05S1 and a Hyundai Hope on Wheels Young Investigator Award. A.B. is a Blood Cancer United Career Development Program Fellow. Murine studies here were performed the UCSF Helen Diller Family Comprehensive Cancer Center Preclinical Therapeutics Core facility, directed by V.S. and supported by NIH award P30CA082103. Molecular dynamics simulations were performed using HPC resources at the Flatiron Institute, a division of the Simons Foundation. A.P.W., D.F., and T.K. are members of the San Francisco Biohub Investigator program.

## Methods

### Deep Learning and physics-based protein design and mutational effect prediction

For structure preparation, chains A and B from CD70 and chain F from CD27 were extracted from PDB 7KX0. For sequence design and design selection, ProteinMPNN (vanilla model) was run on the heteromeric complex 2000 times on interface residues from CRD 2 and 3 of CD27 at 0.1 sampling temperature, with noninterface residues fixed. Mutations at each position as made by ProteinMPNN were compiled into a Rosetta resfile along with the wildtype residue. A Rosetta FastDesign protocol was used to model combinations of mutations as guided by the resfile for design. Outputs were filtered to have less than 6 mutations and shape complementarity and interface stability similar to or better than wild-type, leaving 12 designs. After a round of visual review, 4 designs were selected for experimental testing. For hotspot analysis, Rosetta Flex ddG^54^ analysis was used to identify hotspot interface positions involved using the individual mutations from each of the 4 combinatorial designs selected as described above and modeling them individually as well as mutating the positions to alanines. Positions that disrupt interface stability after mutation to alanine (ΔΔG > 1) were considered hotspots. ΔΔG Scores are reported from averaging across nstruct 35 after 25 backrub steps with the recommended sampling parameters: nstruct = 35, max_minimization_iter = 5000, abs_score_convergence_thresh = 1.0, number_backrub_trials = 3500, backrub_trajectory_stride = 7000

### Tumor Cell Line Culture

Adherent cell lines were grown in DMEM media (Gibco 10564011) supplemented with 10% heat-inactivated fetal bovine serum (FBS) (Gemini) and 100 U/mL penicillin/streptomycin. Non-adherent cell lines were grown in complete RPMI-1640 medium (Gibco 11875093) supplemented with 10% heat-inactivated FBS (Gemini) and 100 U/mL penicillin/streptomycin. All cells were grown at 37 °C with 5% CO2.

### Primary Human T-cell Isolation and Culture

Primary T cells were isolated from LeukoPaks (Stem Cell Technologies, 200-0092) using an EasySep human T cell isolation kit (StemCell, 17951). T-cells were cultured in CTS OpTmizer medium with CTS supplement (Thermo Scientific, A1048501) supplemented with 5% human AB serum (HP1022; Valley Medical), 1% GlutaMAX (Gibco™. Cat # 35050061), 100 U/mL penicillin/streptomycin (Fisher Scientific, 15-140-122), and recombinant interleukin-7 (PeproTech, 200-07) and interleukin-15 (PeproTech, 200-15) at 10 ng/mL.

### Generation of TRAC Knock-in CAR T-cells

After thawing and seeding CD3-isolated T-cells at a concentration of 1 million cells/mL, T cells were stimulated with CD3/CD28 Dynabeads (11131-D; Thermo Fisher Scientific) according to the manufacturer’s instructions (25 μL of beads per 1 million T-cells) for at least 48 hours. T-cells were then isolated from the beads using magnetic separation and electroporated with Cas9-guide RNA complex against *TRAC* and other necessary genes (e.g. *CD70*) as per the nucleofection protocol. T-cells were then recovered in T-cell media without added serum, and, following a 30 minute recovery period, transduced with AAV containing a homology-directed repair template at a selected MOI (e.g. 50,000). After an overnight incubation with AAV, T-cells were counted, washed, and re-suspended in complete T-cell media. Knock-in and knockout efficiency was assessed by flow cytometry at least 72 hours after transduction.

### Generation of Lentivirally Transduced CAR-T cells

CD3-isolated T-cell populations were thawed and seeded at a concentration of at least 1 million cells/mL. Following overnight recovery, if necessary, T-cells were nucleofected with Cas9-guide RNA complex as per the nucleofection protocol. Following overnight recovery after CRISPR editing, T cells were stimulated with CD3/CD28 Dynabeads (11131-D; Thermo Fisher Scientific) for 72 hours. Transduction with CAR lentivirus was performed 1 day after the start of bead stimulation, using approximately 50 microliters of concentrated lentivirus per million T-cells.

Transduction efficiency was assessed by flow cytometry at least 48 hours after the removal of stimulation beads.

### CRISPR-Cas9 Nucleofection

Genetic knockout cells were generated using *in vitro* nucleofection of CRISPR Cas9 ribonuclease protein complexed to sgRNA against the gene of interest. Unless otherwise specified, for each nucleofector well, generally containing 2 million cells, 1.5 μl of each sgRNA (100 µM; Synthego Corporation) and 1.5 μl recombinant Cas9 protein (40 µM; QB3 MacroLab, University of California, Berkeley) was incubated at 37 °C for 15 min to generate a gRNA-ribonuclease complex, which was then nucleofected into the cells of interest using a 4D-Nucleofector (Lonza) with the built-in program DS-137 for cell lines (using Lonza V4XC-2032 kit) and EO-115 for primary T cells (using Lonza V4XP-3032 kit).

### Adeno-Associated Virus (AAV) Production

AAV was produced by transfecting adherent HEK293T cells with CAR-containing HDR template plasmids, along with AAV6 Rep-Cap gene-containing plasmid and E2A, E4, and VA helper gene-containing plasmids using Transfection-grade PEI 25K reagent (Polysciences #23966).

After 72-96 hours, AAV was harvested by concentrating supernatant using PEG-8k at a 1:4 ratio, incubating for 3 hours, and concentrated at 4500 ✕ g for 15 mins. The cell pellet was lysed using rapid freeze-thaw thermal shock and combined with the virus from the supernatant. This combined solution was incubated with 25 U/mL Benzonase (Millipore Sigma #70-664-3) for 1 hour, then centrifuged at 4500 ✕ g. The supernatant was transferred to the top of an iodixanol gradient consisting of layers with 15%, 25%, 40%, and 54% iodixanol at ultracentrifuged at 28,000 rpm overnight in an L8-80M ultracentrifuge (Beckman Coulter) with SW28 rotor (Beckman Coulter). AAV was then extracted from the 40% iodixanol layer and at the 40%/54% iodixanol interface and concentrated in a 50 kDa Amicon centrifugal filter (Millipore Sigma #UFC905024) with buffer exchange into a PBS + 0.001% Tween-20 buffer. AAV titers were obtained using qPCR following a DNase I (NEB #B0303S) and Proteinase K (Qiagen #1114886) digestion to release viral capsid DNA. qPCR was performed using the SsoFast Eva Green Supermix (Bio-Rad #1725201) on a StepOnePlus Real-Time PCR System (Applied Biosystems #4376600) with primers targeting the viral genome. Titers were quantified using a standard curve generated from pure AAV plasmids of known DNA concentration.

### Lentivirus Production

Lenti-X 293T cells (Takara, Cat # 632180) were transfected with CAR expression plasmid and Mirus Bio™ TransIT™-Lenti Transfection Reagent (Cat # MIR6600) as per the manufacturer’s protocol. Transfected Lenti-X 293T cells were cultured for 3 days, with lentivirus being harvested at 48 and 72 hours. Lentivirus was concentrated using Lenti-X Concentrator (Takara Bio, 631232) according to the manufacturer’s protocol.

### Molecular Cloning and DNA Plasmids

Genes encoding the CAR binders were synthesized as gene fragments from Twist Biosciences and cloned into an AAV HDR template or lentiviral expression vector via NEB HiFi assembly (NEB, E2611S). Whole plasmid DNA sequencing was performed to confirm accuracy of the vector. This final construct was then expressed in Stellar Chemically Competent E. coli (Takara Biosciences). DNA was isolated using the QIAGEN Plasmid Plus Midi Kit.

### Flow Cytometry

Immunostaining of cells was performed as per the instructions from the antibody vendor unless stated otherwise. Up to one million cells were resuspended in 50 µl of FACS buffer (PBS + 2% FBS) with 5µL of human Fc Block (Biolegend, 422302) added. Cells were incubated at room temperature for 10 min, then 1-5µL of antibody was added (based on prior titration). Cells were incubated at 4°C for 30 minutes and washed three times with FACS buffer before flow cytometry analysis. For fluorescence compensation, UltraComp eBeads (Invitrogen, 01-2222-42) with 1 µL of antibody added was used. Flow cytometry analysis was performed on the CytoFLEX platform (Beckman Coulter), and data were analyzed using FlowJo_v10.10.0.

### In Vitro Luminescence Cytotoxicity Assay

Target cell lines were engineered to stably express luciferase using lentiviral transduction^31,84^. For each cell line, 50,000 suspension or 10,000 adherent tumor cells were seeded in 50 uL of complete RPMI-1640 media in a white 96 well flat-bottom plate (Greiner Bio-One).

Subsequently, an appropriate number of CAR-T cells were added at various effector:target ratios in 50 uL of complete T-cell media with *n* = 3 replicates per condition. This coculture was incubated at 37°C for 18-24 hours. The next day, 100uL of D-Luciferin (Gold Biotechnology, LUCK-1G) was added to each well to a final concentration of 375 μg/ml, followed by luminescence detection using GloMax Explorer (Promega). The bioluminescence readings were averaged amongst the *n* = 3 technical replicates and the background values were subtracted. Bioluminescence readings were normalized to a cytotoxicity scale of 0-100 using the negative control condition for each E:T ratio.

### Incucyte Live-cell Imaging Cytotoxicity Assay

Target cell lines were engineered to stably express mCherry fluorescent protein using lentiviral transduction. 10,000 target suspension cells were fixed to the bottom of a 96 well flat-bottom plate pre-treated with poly-L Ornithine (Sigma-Aldrich, P4957) for 1 hour in complete RPMI-1640 media. CAR-T cells were added at various effector:target ratios in complete T-cell media with *n* = 3 replicates per condition. The plate was imaged in an Incucyte S3 instrument (Sartorius) and imaged every 4 hours for 72 hours. Data was analyzed on the Incucyte cell-by-cell analysis software, using the mCherry fluorescence as a measure of target cell survival and growth.

### Fluorescent Recombinant Protein Binding Assay

Recombinant trimeric CD70 protein conjugated to FITC or Alexa Fluor 647 ( Acro Biosystems, CDL-HA246) was used to stain 100,000 CAR-T cells at a concentration of 1 μg/mL for 1 hour at 4°C, followed by an additional antibody stain for cell-specific markers following the flow cytometry protocol. Binding strength was assessed by median fluorescence intensity (MFI) of the fluorophore conjugated to CD70 for CAR-positive cells by flow cytometry. MFI was normalized to the CD27_WT_ CAR value for each experiment.

### Cytokine Production Assay

200,000 target tumor cells were plated in each well of a 96 well flat-bottom plate in complete RPMI-1640 media. An appropriate number of CAR-T cells for each condition in complete T-cell media were added to each well and co-cultured at 37°C for 6 hours. At 1 hour of co-culture, protein transport inhibitors Brefeldin A (Biolegend, 420601) and/or Monensin (Biolegend, 420701) was added to each well at the manufacturer’s recommended 1X final concentration and the plate was returned to the 37°C incubator. Following co-culture, the plate was washed in FACS buffer, stained with a fixable viability dye and antibodies for cell surface markers, and incubated for 30 mins (see flow cytometry protocol). Samples were then fixed using an intracellular paraformaldehyde fixation buffer (Biolegend, 420801) for 1 hour at RT in the dark, washed with FACS buffer, then washed with permeabilization buffer (Biolegend, 421002).

Samples were stained using antibodies against intracellular targets, including cytokines of interest, for 30 minutes at 4°C, then washed with permeabilization buffer and analyzed by flow cytometry.

### Phospho-ERK 1/2 stimulation Assay

50,000 target cells were added to each well of a 96 well round-bottom plate in complete RPMI-1640 media. An appropriate number of CAR-Ts for each E:T ratio in complete T-cell media were added at T = 0, 30, 45, 50, and 55 minutes with light mixing and co-cultured at 37°C. At T = 60 minutes, warm paraformaldehyde fixation buffer (Biolegend, 420801) was added to all wells and incubated at 37°C for 15 mins. All samples were washed with FACS buffer, then permeabilized with ice-cold True-Phos Perm Buffer (Biolegend, 425401) for 60 mins at -20°C. Samples were washed with FACS buffer, then stained with antibodies for extracellular and intracellular markers, including anti-ERK1/2 Phospho (Thr202/Tyr204), following the flow cytometry protocol and analyzed on a flow cytometer.

### Murine Cell Line Xenograft CAR-T Studies

Murine studies were conducted in accordance with UCSF Institutional Animal Care and Use Committee protocol and guidelines. NSG (NOD.Cg-*Prkdc^scid^ Il2rg^tm1Wjl^*/SzJ, strain #:005557, Jackson Laboratories) male or female mice, 8-10 weeks old, were bred at the UCSF Breeding Core Facility. For hematological cancer cell lines, 1 million cells were injected IV via the tail vein. For solid tumors, 5 million cells were mixed with matrigel (Corning, 354234) and injected SC into the flank. Several days following tumor injection, mice were transfused IV with CAR-T cells or untransduced T cells. Bioluminescence imaging (Perkin Elmer In Vivo Imaging System, Caliper Life Sciences) was performed to assess tumor burden in disseminated hematological tumor models and caliper measurements of flank tumors were performed to assess tumor size in solid tumor models. Peripheral blood draws were performed to assess tumor burden and CAR-T expansion at a predetermined frequency. Survival end point of the study was determined by signs of symptomatic illness in the animals and euthanasia humane endpoints as consistent with the IACUC protocol.

### Single-Cell RNA Sequencing

Raw sequencing data was processed using Cell Ranger (v8.0.0). FASTQ files were aligned to GRCh38 and quantified using Cell Ranger count with default settings. Raw counts were normalized to 10,000 counts and then log transformed after incrementing by 1 using Seurat (v5.5.1)^85^. Clustering and visualization was performed using Monocle3 (v1.4.27)^86^. CD4 and CD8 single T-cells were selected for further analysis based on single cell CD4 and CD8 expression, respectively. Reads associated with unfiltered, CD4, and CD8 cells were aggregated for pseudobulk analysis within each cell type population. Differential expression was performed using edgeR (v4.8.2)^87^.

### Molecular Dynamics Simulations

Extensive all-atom molecular dynamics (MD) simulations^88^ were performed on both the wildtype (WT) and N88A mutant trimeric complexes derived from the crystal structure (PDB ID: 7KX0)^53^. The complexes in the study comprised the complete CD27-CD70 hexameric assembly (the inner CD70 trimer with chains A, B, and C, and the outer CD27 trimer with chains D, E, and F), with all crystallographically resolved N-linked glycans retained, to characterize mutation effects across all three equivalent binding interfaces where each outer CD27 monomer contacts two inner CD70 monomers (CD70-CD27 pairs of BC-D, AC-E, and AB-F) under physiologically relevant conditions. Each complex was solvated in a rectangular periodic box of explicit TIP3P water molecules^89^, with a minimum distance of 12 Å between the solute and box edges. The complexes were then neutralized and brought to a physiological ionic strength of 150 mM by addition of K^+^ and Cl^-^ ions, followed by parameterization employing the CHARMM36m force field^90^. Prior to production dynamics, complexes were subjected to energy minimization using the L-BFGS algorithm with a convergence tolerance of 100 kJ mol^-1^ nm^-1^ to remove steric clashes and unfavorable contacts. Production simulations were performed employing the OpenMM 8.0 MD engine^91^. The temperature was maintained at 300 K using the Langevin integrator, and a friction coefficient of 1 ps^-1^ was employed to provide efficient thermostatting while minimizing perturbations to the system dynamics. The equations of motion were integrated with a 2 fs timestep, enabled by constraining all covalent bonds involving hydrogen atoms using the SHAKE algorithm^91^. Long-range electrostatic interactions were computed using the particle mesh Ewald (PME) method^92^ with a cutoff of 1.0 nm. Three independent replicates, each with a simulation length of 2.25 μs, were run for both WT and N88A complexes, yielding a cumulative simulation time of 6.75 μs per complex. Trajectory coordinates were saved every 200 ps, and system energetics and temperature were monitored throughout each simulation to verify equilibration and stability.

### Biolayer Interferometry

In a 96 well flat-bottom plate, Streptavidin biosensors (Sartorius, 18-5019) were pre-equilibrated in an assay buffer of 1X PBS, 0.05% Tween-20, 0.2% BSA) for at least 15 minutes. A predetermined concentration (e.g. 25nM) of the biotinylated target protein (e.g. CD70) diluted in the assay buffer was added to one well per condition in a 384 well flat-bottom plate. The binding partner ligand (e.g. CD27_var_ recombinant protein) was serially diluted three-fold from a predetermined maximum concentration at least 10-fold higher than the expected dissociation constant in a blocking buffer (1X PBS, 0.05% Tween-20, 0.2% BSA, 10μM biotin). Each concentration was added to a separate row of the 384 well plate. Additional wells were filled with assay buffer and blocking buffer for each ligand concentration added. On an Octet RED384 (Sartorius) BLI instrument, SA biosensors were immersed in assay buffer for 90 seconds to reach the first baseline, then loaded with biotinylated target protein for 500 seconds, placed into blocking buffer for 180 seconds for a second baseline, placed into the appropriate concentration of ligand protein for 600 seconds to measure the association rate, then re-placed into blocking buffer for 1500 seconds to measure the dissociation rate. This experiment was performed at 30°C at a plate mixing rate of 1000 rpm. Data was analyzed on the Octet Analysis Studio Software, subtracting reference sensors and reference wells and aligning samples to the average of the baseline step with Savitsky-Golay noise filtering. Kinetic analysis was performed using a 1:1 Global fit model to determine the association and dissociation rate constants and overall dissociation constant (K_d_).

### Cell Doublet Avidity Assay

Target tumor cells engineered to express mCherry were plated at in a 96 well U-bottom plate, with 2.5×10^5^, 3.75×10^5^, or 4.5×10^5^ cells in each well, corresponding to an E:T ratio of 1:1, 1:3, or 1:9, respectively. A total of 2.5×10^5^, 1.25×10^5^, or 5.0×10^4^ CAR T-cells, respectively, were stained with antibody against a CAR transduction marker and a T-cell marker for 30 mins at 4°C, washed in FACS buffer, then added to each well to achieve the appropriate E:T ratio, with a total of 5.0×10^5^ cells per well. Following a 60 minute co-culture at 37°C, the plate was loaded onto a flow cytometer without washing and doublets were calculated based on events that were positive for both mCherry and a T cell/CAR marker, indicating a CAR-T cell attached to a tumor cell.

### Trogocytosis Assay

Tumor cells and CAR T-cells were plated in a 96 well flat-bottom plate at a 1:2 E:T ratio and co-incubated at 37°C. At selected time points (e.g. 1, 2.5, and 4 hours), approximately 25% of the co-culture volume was removed from the plate and flow cytometry was run on the sample after staining for cell-specific markers and target antigens. Median fluorescence intensity was measured on CAR T-cells and graphed over time.

### Total Internal Reflection Fluorescence Microscopy

CAR-Ts were added directly to planar supported lipid bilayers (SLBs), made from small unilamellar vesicles and MOPS buffer as previously described. These SLBs contained FITC-labeled trimeric CD70 (Acrobiosystems) and cells were allowed to settle for 20 minutes prior to imaging. Engaged cells were located and identified using reflection interference contrast microscopy (RICM). Images were acquired using RICM and TIRF microscopy for all relevant fluorescent channels. When applicable, CD27 and CD45 were labeled with 10 nM anti-CD27 and anti-CD45 antibody, respectively. Cells were imaged with a TIRF microscope (Eclipse Ti, Nikon), 60x objective (Apo TIRF, 1.49NA, oil, Nikon) and EMCCD camera (iXON Ultra, Andor). RICM images were flattened using a background (no cells or GPMV) image, then outlines of SLB-bound cells or GPMVs were identified. Fluorescence inside and outside these ROI outlines was measured and averaged for each fluorescent channel. The outside background measurement was subtracted from the average signal inside of the footprint using an automated custom macro in FIJI.

### Off-target Protein Binding Assay

This assay utilized the Retrogenix^®^ Cell Microarray screening platform. HEK293 cells were reverse-transfected to overexpress 6108 unique human plasma membrane and membrane-tethered secreted proteins, including the CD70 target antigen. Each construct also included ZsGreen1 as a positive control for transfection. Initial screens tested the optimal concentration of CD27_N88A_ for signal to noise ratio using CD70, EGFR, and untransfected HEK293 cells negative control. Binder specificity screens using the full library were performed by spotting constructs in duplicate onto slides, and reverse-transfecting and fixing HEK293 cells. The slides were incubated with CD27_N88A_ followed by detection of binding using a secondary fluorescent antibody. This screen was performed in duplicate and all identified interactions were repeated in a confirmatory screen, performed on both fixed and live cells, with CD27_N88A_ or 4-1BB negative control. Standard Retrogenix^®^ thresholds of signal to noise ratios are used to assess true- vs. false-positive binding.

### HSPC Colony Forming Assay

CD34+ cells were isolated from healthy donor GM-CSF-mobilized peripheral blood (Stemcell Technologies EasySep Human CD34 Positive Selection Kit II) and co-incubated with CAR-T cells in IMDM media with added 2% FBS and penicillin/streptomycin at a 5:1 E:T ratio for 5 hrs in triplicate in a U-bottom 96-well plate. 1000 CD34+ cell-containing co-culture was then transferred to a total volume of 1.1 mL of methylcellulose-based medium (MethoCult H4434 Classic, StemCell Technologies) per condition in a 35 mm dish and placed in a 37°C cell culture incubator for 14 days. Colonies were counted and classified as granulocyte-erythroid-macrophage-megakaryocyte colony forming unit (CFU-GEMM), granulocyte-macrophage colony forming unit (CFU-GM), burst-forming unit-erythroid (BFU-E), or erythroid colony forming unit (CFU-E). Brightfield images were acquired with a Keyence BZ-X800 microscope at 4x magnification, and the number of each colony type was tallied and averaged from triplicate conditions.

### Code Availability

Code for Rosetta and ProteinMPNN modeling is available at: https://github.com/radhikadalal99/N88A_fastdesign_protocol

Link for molecular dynamics simulation trajectories are available at: https://zenodo.org/records/22000829

### Data Availability

Single-cell RNA-seq has been deposited at the Gene Expression Omnibus; accession number is pending and will be provided at time of publication.

### Statistical Analyses

All hypotheses were tested at an α=0.05 level unless otherwise indicated. Data was analyzed using GraphPad Prism 10, and specific statistical tests used for each experiment have been indicated within the main text or figure legends.

