## Supplemental Figures and Tables for "Deep Learning-based Modeling Enhances Efficacy of Natural Ligand CAR Binders Targeting CD70"

### Figure S1

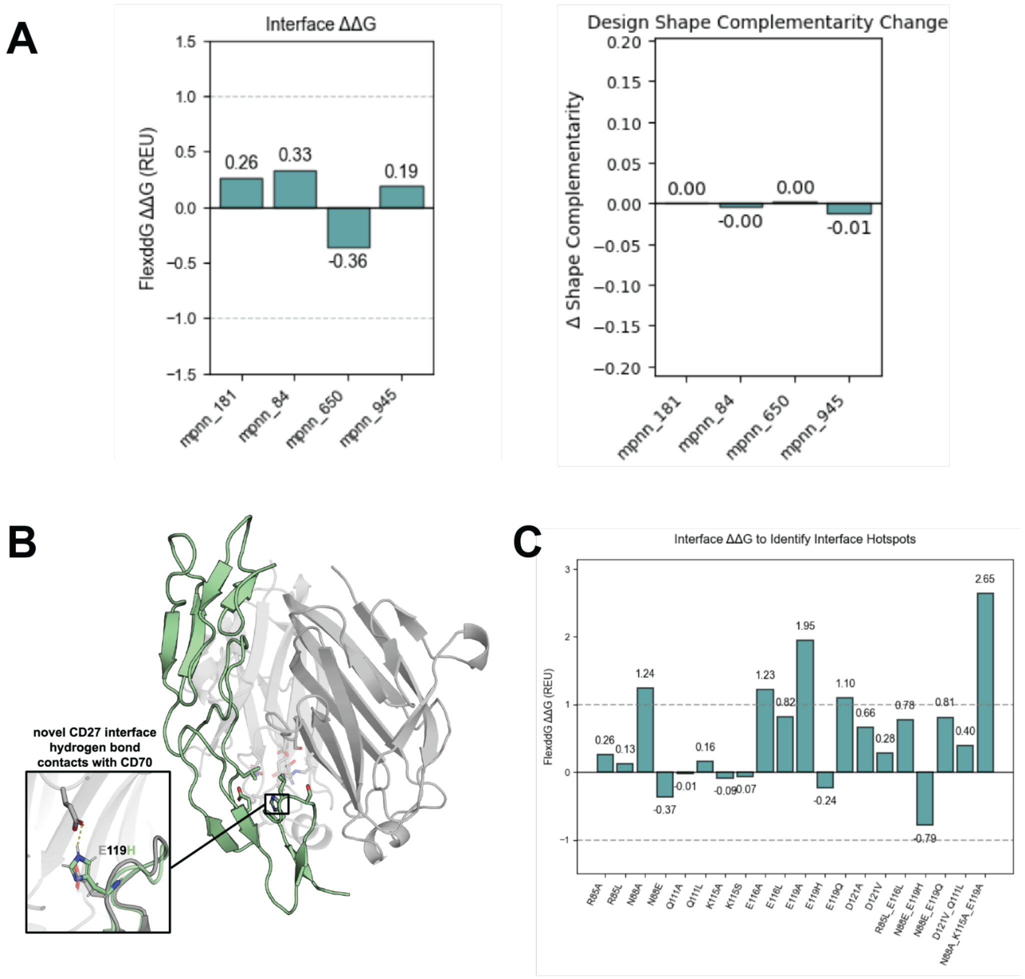


**Figure S1. Rosetta interface metrics for designed variants. A-B.** Combinatorial designs are predicted to maintain similar interface binding energies (ΔΔG between -1 and 1 Rosetta Energy Units, REU, *left*) and similar shape interface complementarity (*right*) relative to the wild-type structure while introducing new predicted contacts. **C.** Analysis of predicted effects of designed individual and combinatorial mutations on interface stability using Rosetta flex ddG. “Hot spots” are typically defined as positions with ΔΔG > 1 REUs (dotted line) when mutated.

### Figure S2

**A**

**Generation of TRAC Knock-in CAR T-cells**

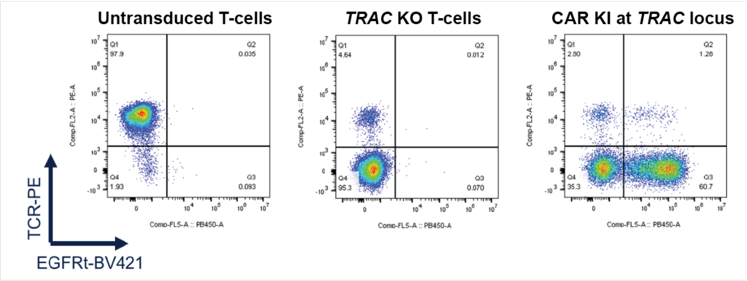


**B**

**Relative Transduction Marker Expression Across CD27 CAR T-cell Variants**

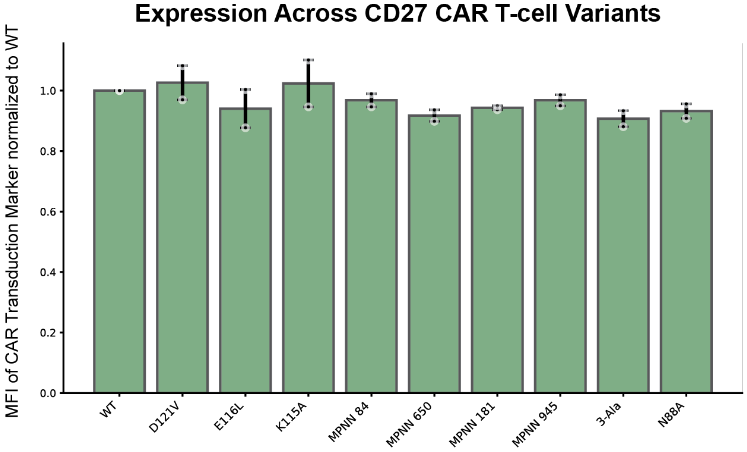


**Figure S2. $CD27_{var}$ CAR-Ts express effectively and uniformly in the *TRAC* locus. A.** Representative T-cell flow cytometry plots for *TRAC* knockout, indicated by TCR surface expression, and CAR knock-in, as measured by truncated EGFR ("EGFRt") transduction marker expression. **B.** Median fluorescence intensity of EGFR transduction marker in CAR expression cassette across CD27 CAR variants, normalized to the MFI of $CD27_{WT}$ in *n*=2 healthy donors. Error bars represent S.D.

### Figure S3

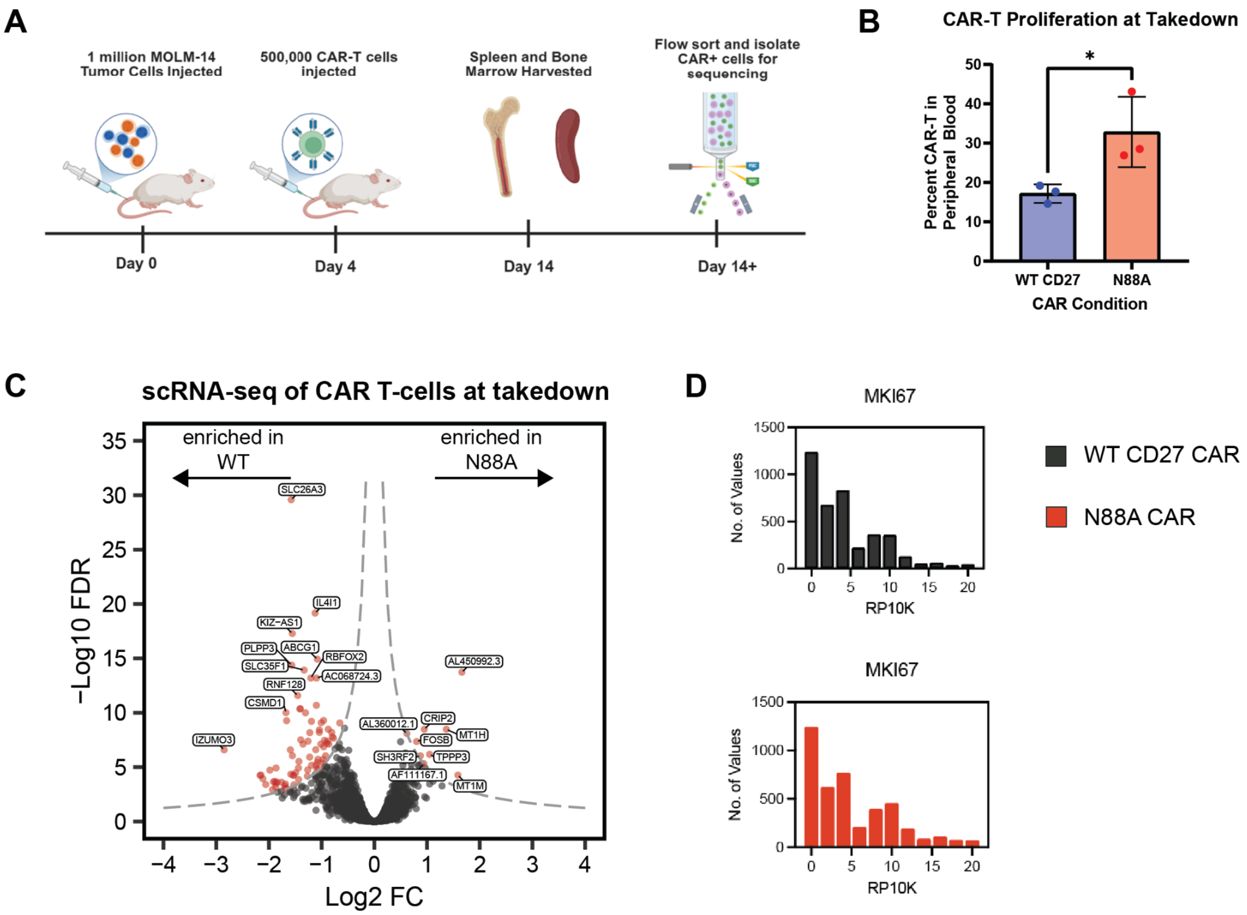


**Figure S3. CD27$_{N88A}$ CAR-Ts display altered transcriptomics compared to CD27$_{WT}$ in an *in vivo* model. A.** Cartoon overview timeline of scRNA-seq experiment to compare transcriptional states of CD27$_{WT}$ versus CD27$_{N88A}$. **B.** Flow cytometry-based measurement of CAR-T proliferation in each treatment group as a percent of all live non-RBC events during FACS sort of processed mouse spleen tissue. Error bars represent S.D., *p*-value by student's *t*-test **C.** Volcano plot of differentially transcribed genes between CD27$_{WT}$ and CD27$_{N88A}$ samples via pseudobulk analysis. $n$ = 3 mice per study arm. **D.** Histogram of transcript levels of proliferation marker MKI67 in CD4+ populations of CAR T-cells, average of $n$ = 3 replicates per group. *p<0.05; **p<0.01; ***p<0.005; ****p<0.001.

### Figure S4

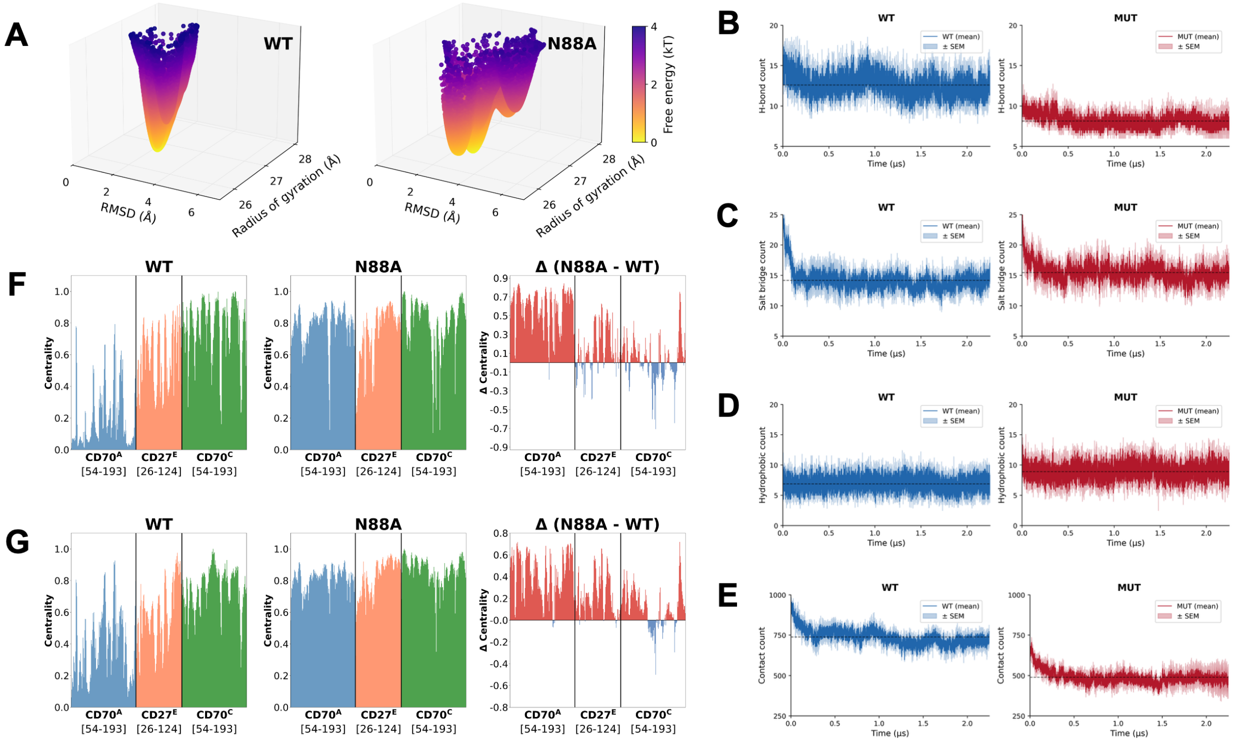


**Figure S4. Molecular dynamics-based contributions of molecular interactions to CD27-CD70 complex free energies. A.** Three-dimensional representation of the free energy landscape showing the energy surface topology. WT (left) features a single deep funnel characteristic of a stable bound complex. N88A (right) exhibits a rugged landscape with multiple local minima separated by low barriers (<1 kBT), facilitating transitions between conformational substates. **B.** Time-resolved H-bond profiles for the WT and N88A CD27-CD70 complexes. The mean interaction count is shown with the standard error of the mean indicated by the shaded region, calculated across all three trimeric CD27-CD70 interfaces and three independent 2.25 μs replicates. Dashed horizontal lines indicate the trajectory mean for each complex. **C.** Time-resolved salt bridge profiles for the WT and N88A ("MUT") CD27-CD70 complexes. **D.** Time-resolved hydrophobic contact profiles for the WT and N88A CD27-CD70 complexes. The mean interaction count is shown with the standard error of the mean indicated by the shaded region, calculated across all three trimeric CD27-CD70 interfaces and three independent 2.25 μs replicates. Dashed horizontal lines indicate the trajectory mean for each complex. **E.** Time-resolved non-bonded contact profiles for the WT and N88A CD27-CD70 complexes, calculated using a 4.5 Å cutoff for total atomic contacts. **F.** Eigenvector centrality profiles derived from linear mutual information correlation networks for the WT and N88A CD27-CD70 functional unit. The left panel shows the WT centrality, the center panel shows the N88A centrality, and the right panel shows the difference (∆ = N88A– WT), where positive values (red) indicate increased centrality and negative values (blue) indicate decreased centrality upon mutation. **G.** Eigenvector centrality profiles derived from mutual information correlation networks for the WT and N88A CD27-CD70 functional unit.

### Figure S5

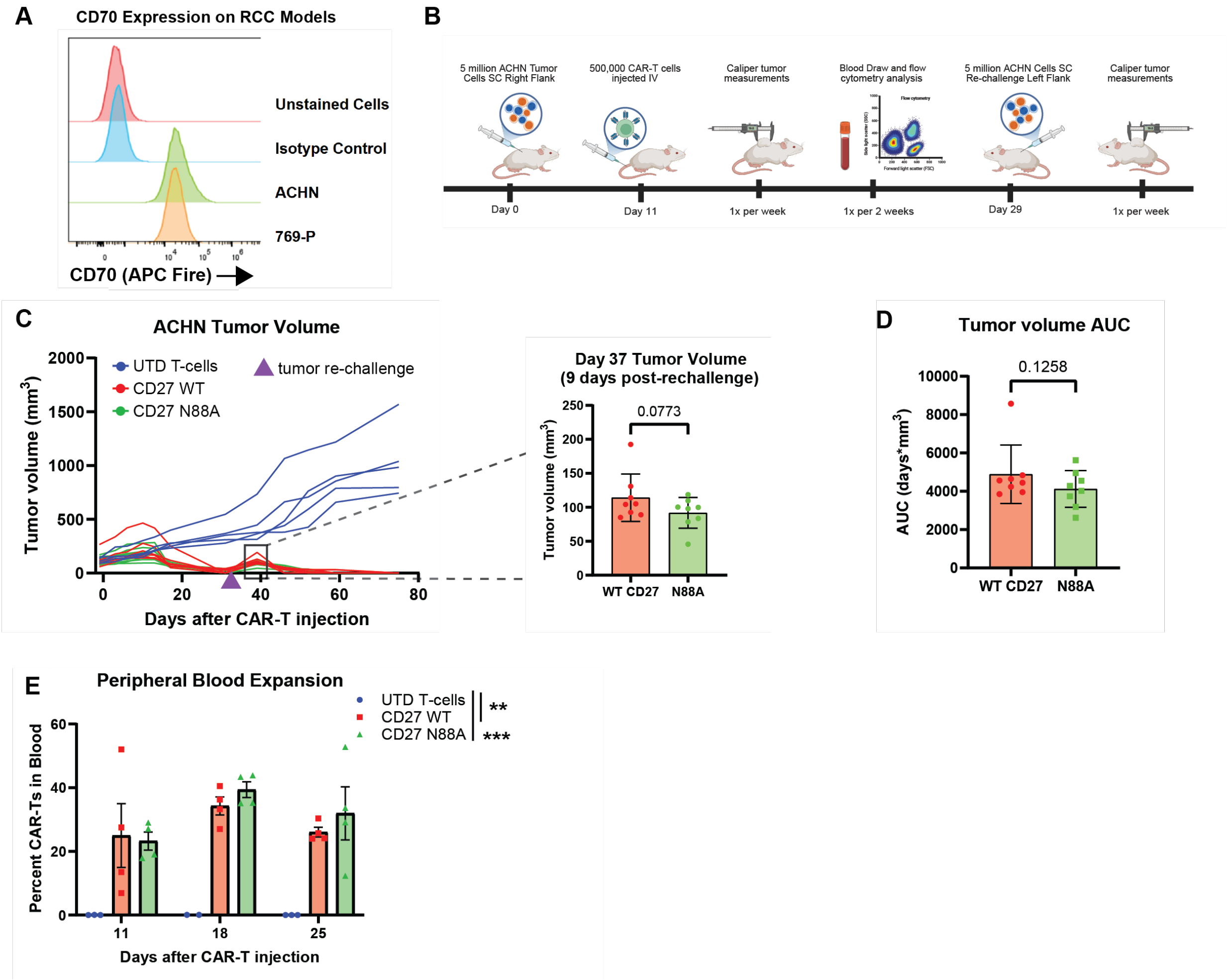


**Figure S5. CD27$_{var}$ CAR-Ts are highly efficacious in tumor control of ACHN solid tumor renal cell carcinoma *in vivo* model and NOMO-1 AML *in vivo* model. A.** Expression of CD70, as measured by flow cytometry, on renal cell carcinoma cell lines ACHN and 769-P. **B.** Overview timeline of *in vivo* CAR-T mouse xenograft re-challenge experiment with ACHN cell line. Tumor rechallenge was performed on day 31. SC = subcutaneous. **C.** *Left:* quantification of tumor burden following bioluminescence imaging of mice at select time points after CAR-T injection. *Right, inset:* Comparative tumor volume in CAR T-treated mice on day 37, 9 days post-rechallenge. p-value by one-tailed t-test. **D.** Area-under-the-curve comparison of tumor volume vs time for all individual mice in CAR T-treated mice, including initial tumor challenge and re-challenge. **E.** Flow cytometry-based measurement of CAR-T proliferation in each treatment group, as a percent of all live, non-RBC, singlet events from weekly or biweekly blood draws. n=3 blood draws per group at each time point. *p*-value by 2-way ANOVA with Tukey's correction. **F.** Overview timeline of *in vivo* CAR-T mouse xenograft re-challenge experiment with NOMO-1 AML cell line. Tumor rechallenge was performed on day 31. **G.** Kaplan-Meier survival curve of mice by treatment group. Mice were sacrificed at clinical humane endpoint due to tumor burden following the recommendations in the approved IACUC protocol. P-value by log-rank. **H.** *Left:* Quantification of tumor burden following bioluminescence imaging of mice at select time points after CAR-T injection. *Right, inset:* Comparative tumor volume in CAR T-treated mice on day 39, 8 days post-rechallenge. *p*-value by unpaired *t*-test. **I.** Flow cytometry-based measurement of CAR-T proliferation in each treatment group, as a percent of all live, non-RBC, singlet events from weekly or biweekly blood draws. *n*=3 blood draws per group at each time point. *p*-value by unpaired t-test. *p<0.05, **p<0.01, ***p<0.005.

### Figure S6

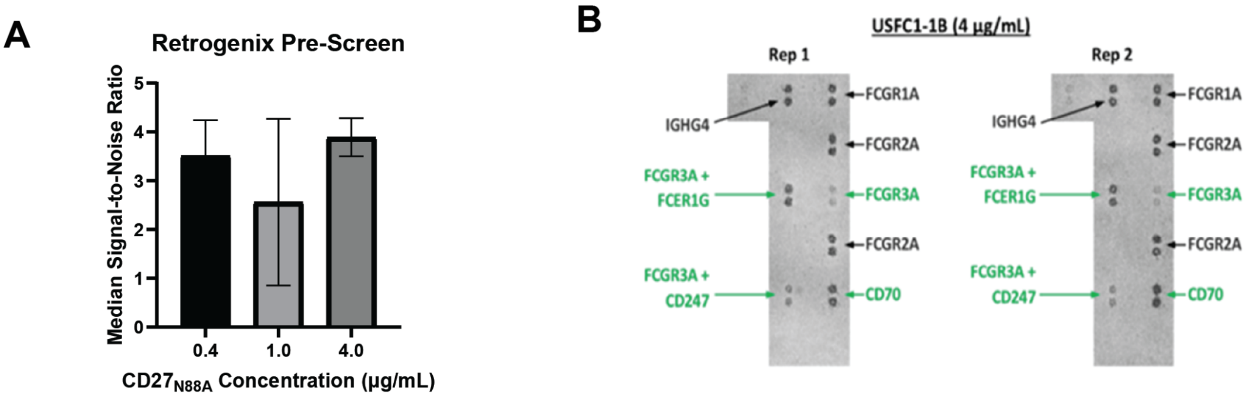


**Figure S6. CD27$_{N88A}$ recombinant protein shows no significant off-target binding in the Retrogenix® screening platform. A.** Assessment of median signal-to-noise ratio of CD27$_{N88A}$ binding to CD70 at multiple binder concentrations, identifying 4 μg/mL as the optimal assay concentration. Error bars represent data range, n=3 replicates. **B.** Representative images of Retrogenix® off-target confirmation screening, evaluating recombinant CD27 N88A protein binding to HEK293 cells expressing individual cell-surface proteins identified in the library screen; n=2 technical replicates per target protein and n=2 experimental replicates per binder tested.

| # | Lost | Gained |
|---|---|---|
| 1 | S89(E)–N88(E) | S89(F)–N88(F) |
| 2 | N88(E)–C87(E) | S89(D)–N88(D) |
| 3 | S89(D)–N88(D) | S89(E)–N88(E) |
| 4 | S89(F)–N88(F) | N88(F)–C87(F) |
| 5 | N88(D)–C87(D) | N88(D)–C87(D) |
| 6 | N88(F)–C87(F) | N88(E)–C87(E) |
| 7 | N88(F)–S178(A) | N88(E)–T118(E) |
| 8 | N88(D)–S178(B) | N88(F)–T118(F) |
| 9 | N88(E)–T118(E) | N88(D)–T118(D) |
| 10 | N88(E)–S178(C) | N88(F)–H86(F) |

**Table S1. Hydrogen bond changes upon N88A mutation at the CD27-CD70 interface.**

| # | Lost | Gained | Increased | Decreased |
|---|---|---|---|---|
| 1 | H124(C)–D121(D) | R83(C)–E82(E) | R144(B)–D182(A) | R157(B)–E59(A) |
| 2 | – | R85(F)–E119(F) | R85(E)–D182(C) | R144(C)–D114(D) |
| 3 | – | R113(F)–E183(A) | R144(B)–E183(A) | R144(B)–D114(F) |
| 4 | – | K115(F)–E119(F) | R85(D)–D182(B) | R85(E)–E116(E) |
| 5 | – | H124(C)–D114(D) | R83(A)–E82(F) | R113(F)–D121(F) |
| 6 | – | R157(A)–D182(C) | R144(C)–D182(B) | R144(A)–D182(C) |
| 7 | – | H123(A)–D121(E) | R113(E)–D121(E) | R144(A)–D114(E) |
| 8 | – | – | R144(C)–E116(D) | R85(F)–D182(A) |
| 9 | – | – | R85(D)–E183(B) | R144(A)–E183(C) |
| 10 | – | – | R113(D)–E119(D) | R113(D)–D121(D) |

**Table S2. Salt bridge changes upon N88A mutation at the CD27-CD70 interface.**

| # | Lost | Gained | Increased | Decreased |
|---|---|---|---|---|
| 1 | – | A113(B)–N88(D) | A113(B)–L176(B) | L112(B)–L175(B) |
| 2 | – | A113(A)–N88(F) | A102(E)–L92(E) | L112(C)–L175(C) |
| 3 | – | A113(C)–N88(E) | A113(A)–L176(A) | A113(C)–L176(C) |
| 4 | – | N88(F)–I114(A) | A102(D)–L92(D) | A113(A)–I114(A) |
| 5 | – | N88(D)–I114(B) | A105(E)–V93(E) | A113(B)–I114(B) |
| 6 | – | N88(E)–I114(C) | – | A105(F)–V93(F) |
| 7 | – | P122(D)–W110(D) | – | A105(D)–V93(D) |
| 8 | – | – | – | L175(A)–L112(A) |
| 9 | – | – | – | A105(F)–L91(F) |

**Table S3. Hydrophobic contact changes upon N88A mutation at the CD27-CD70 interface.**

| # | Lost | Gained |
|---|---|---|
| 1 | N88(E)–S89(E) | N88(F)–S89(F) |
| 2 | N88(D)–S89(D) | N88(D)–S89(D) |
| 3 | N88(F)–S89(F) | N88(E)–S89(E) |
| 4 | N88(E)–C87(E) | N88(F)–C87(F) |
| 5 | N88(F)–C87(F) | N88(E)–C87(E) |
| 6 | N88(D)–C87(D) | N88(D)–C87(D) |
| 7 | A113(C)–N88(E) | N88(F)–I114(A) |
| 8 | A113(B)–N88(D) | A113(B)–N88(D) |
| 9 | N88(E)–E119(E) | N88(D)–I114(B) |
| 10 | N88(E)–I114(C) | N88(E)–I114(C) |

**Table S4. Non-bonded contact changes upon N88A mutation at the CD27-CD70 interface.**
